# Functional connectivity between the cerebellum and somatosensory areas implements the attenuation of self-generated touch

**DOI:** 10.1101/830646

**Authors:** Konstantina Kilteni, H. Henrik Ehrsson

## Abstract

Since the early 1970s, numerous behavioral studies have shown that self-generated touch feels less intense than the same touch applied externally. Computational motor control theories have suggested that cerebellar internal models predict the somatosensory consequences of our movements and that these predictions attenuate the perception of the actual touch. Despite this influential theoretical framework, little is known about the neural basis of this predictive attenuation. This is due to the limited number of neuroimaging studies, the presence of conflicting results about the role and the location of cerebellar activity, and the lack of behavioral measures accompanying the neural findings. Here, we combined psychophysics with functional magnetic resonance imaging to detect the neural processes underlying somatosensory attenuation in male and female healthy human participants. Activity in bilateral secondary somatosensory areas was attenuated when the touch was presented during a self-generated movement (self-generated touch) than in the absence of movement (external touch). An additional attenuation effect was observed in the cerebellum that is ipsilateral to the passive limb receiving the touch. Importantly, we further found that the degree of functional connectivity between the ipsilateral cerebellum and the contralateral primary and bilateral secondary somatosensory areas was linearly and positively related to the degree of behaviorally assessed attenuation; that is, the more participants perceptually attenuated their self-generated touches, the stronger this corticocerebellar coupling. Collectively, these results suggest that the ipsilateral cerebellum is fundamental in predicting self-generated touch and that this structure implements somatosensory attenuation via its functional connectivity with somatosensory areas.

**Significance statement:** When we touch our hand with the other, the resulting sensation feels less intense than when another person or a machine touches our hand with the same intensity. Early computational motor control theories have proposed that the cerebellum predicts and cancels the sensory consequences of our movements; however, the neural correlates of this cancelation remain unknown. By means of functional magnetic resonance imaging, we show that the more participants attenuate the perception of their self-generated touch, the stronger the functional connectivity between the cerebellum and the somatosensory cortical areas. This provides conclusive evidence about the role of the cerebellum in predicting the sensory feedback of our movements and in attenuating the associated percepts via its connections to early somatosensory areas.

## Introduction

Imagine a situation where your brain cannot differentiate the sensory signals that your body produces from signals that originate from events, objects and actions produced by others in the surrounding environment. In that bizarre situation, the world would appear to constantly move each time you perform a saccade or change your gaze direction, you would continuously wonder whether somebody is talking to you each time you speak, and you would relentlessly tickle yourself each time you touched your own body. One of the strategies the brain uses to avoid such situations is to suppress the perception of self-generated information and thus to magnify its distinction from externally generated input; consequently, self-produced signals feel less intense than signals of identical intensity that are due to external causes (Blakemore et al., 2000; Bays and Wolpert, 2008). This is the classic perceptual phenomenon of sensory attenuation.

In the somatosensory domain, several behavioral studies have shown that the sensations produced by one of our hands voluntarily touching the other hand are systematically attenuated. For example, participants rate a self-generated tactile stimulus on their hand as less intense (and less ticklish) than an external stimulus of the same intensity and frequency (Weiskrantz et al., 1971; Blakemore et al., 1999a). Similarly, in a force discrimination task, participants judge an external tap on their finger to be stronger than a self-induced tap of the exact same intensity (Bays et al., 2005; Kilteni et al., 2019). Moreover, in the classic force-matching task where participants are asked to reproduce the force they just felt on their finger pad, they produce stronger forces than the ones required, which indicates that the self-produced forces feel weaker (Shergill et al., 2003; Wolpe et al., 2016; Kilteni and Ehrsson, 2017a, 2017b).

Computational theories of motor control have suggested that sensory attenuation is a perceptual correlate of the brain’s machinery for motor control. Specifically, it has been theorized that our brain uses internal forward models – probably implemented in the cerebellum (Miall and Wolpert, 1996; Wolpert et al., 1998; Shadmehr and Krakauer, 2008; Shadmehr et al., 2010; Therrien and Bastian, 2018) – to predict the sensory consequences of our actions using the information from the motor command (efference copy) (Kawato, 1999; Bays and Wolpert, 2007; Franklin and Wolpert, 2011). The predictions of these models are necessary to compensate for the intrinsic delays and noise in our sensory system, thus enabling efficient online motor control (Kawato, 1999; Davidson and Wolpert, 2005; Shadmehr and Krakauer, 2008). In addition, these predictions are used to ‘cancel’ the self-induced reafferent input and thus to effectively distinguish it from input produced by external causes (Wolpert and Flanagan, 2001). Consequently, self-generated sensory information is attenuated because it has been predicted by the internal forward models (Blakemore et al., 2000; Frith et al., 2000).

What is the neural basis of somatosensory attenuation? In contrast to the plethora of behavioral paradigms, neuroimaging studies of somatosensory attenuation have been scarce and have provided contradictory results about the brain correlates of the phenomenon. In their seminal study, Blakemore and colleagues (1998) observed reduced activity in the bilateral secondary somatosensory cortex and in the cerebellum contralateral to the passive limb receiving the touch when the touch was presented in the context of a voluntary movement (self-generated touch) compared to when the participants remained motionless (touch generated by an external cause). These observations were based on a very small sample size (six volunteers) and using a fixed-effect analysis. The authors proposed that the reduced cerebellar activity reflects the discrepancy between the predicted and the actual touch – the prediction error – that is at minimum during self-generated touch. In contrast, Shergill and colleagues (2013) observed an increase, rather than a decrease, in the activity of the contralateral cerebellum when directly contrasting a condition involving self-generated touch versus a condition involving external touch, contradicting the proposal of Blakemore. In a subsequent study, Blakemore et al. (2001) found increased cerebellar blood flow with increasing delays between the movement of the active hand and the resulting touch on the passive hand, providing evidence once again that cerebellar activity reflects the prediction error. In contradiction to Blakemore, Shergill and colleagues (2013) failed to observe changes in cerebellar activation when a delay was introduced between the pressing movement of the active hand and the resulting touch on the passive hand, again calling into question the contribution and role of cerebellar activity in somatosensory attenuation.

An additional observation that remains unclear concerns the site of cerebellar activity detected by the previous neuroimaging studies. Given that the somatosensory attenuation is observed on the passive limb that is receiving the touch (Shergill et al., 2003; Bays et al., 2005) and that the somatotopic representations in the cerebellum are mainly ipsilateral (Grodd et al., 2001; Manni and Petrosini, 2004) and the corticocerebellar connections contralateral (Buckner et al., 2011), it is puzzling why the earlier studies observed activations in the cerebellar lobe contralateral to the passive hand. According to the framework of internal models, the predictions attenuating the actual touch should be specific to the passive limb. For example, when we move our right hand to touch the left hand, the brain predicts tactile input on the left hand given the motor command sent to the muscles of the right hand and the proximity between the hands (Bays and Wolpert, 2008; Kilteni and Ehrsson, 2017b, 2017a; Kilteni et al., 2018). Therefore, one would expect that cerebellar activity related to somatosensory predictions or prediction errors concerning the hand receiving the touches should be encoded in the cerebellar hemisphere that is ipsilateral and not contralateral to that hand. However, until now, evidence for such ipsilateral cerebellar responses is lacking, which is problematic because contralateral activation fits neither with the sensorimotor account of internal models nor with human neuroanatomy.

Finally, none of the abovementioned studies included a behavioral assessment of somatosensory attenuation. This is a critical limitation in any study that aims to isolate the neural processes that are specific to sensory attenuation. Although the abovementioned studies revealed a different cerebellar pattern between self-generated and externally generated touch conditions, this does not necessarily mean that the cerebellum is genuinely involved in the predictive attenuation of self-generated touch because no relationship with the behaviorally registered attenuation has been established. Indeed, if the cerebellum is involved in predictive attenuation, one would expect increased cerebellar *interactions* with somatosensory areas for individuals who show stronger behavioral attenuation, indicating that the flow of information between those areas reflects the extent to which participants perceive their touch as weaker than external touch. Nevertheless, to our knowledge, there is no study assessing somatosensory attenuation at both the neural and behavioral levels; therefore, the cerebellar contribution and the neural basis of the phenomenon remain unknown.

To address the abovementioned issues, here we combined functional magnetic resonance imaging (fMRI) with a force-matching psychophysics task and utilized a larger sample of participants than those used in earlier studies. In addition to merely contrasting self-generated touch and externally generated touch, we further took advantage of previous observations that not all self-generated touches are attenuated to the same extent but mainly those that correspond to *direct* self-touch where the two body parts in question are perceived to be in physical contact (Kilteni and Ehrsson, 2017b). For example, in the force-matching task, when the participants reproduce the externally generated forces by moving a joystick that controls the force output on their finger instead of directly pressing their index finger against their other finger, they show no attenuation of their self-generated forces (Shergill et al., 2003; Wolpe et al., 2016; Kilteni and Ehrsson, 2017a). Similarly, if the participants reproduce the externally generated forces by pressing their finger against their other finger but a distance of 15 cm or farther has been introduced between their two hands in the horizontal plane, the attenuation is significantly decreased compared to when the hands are placed close, with one index finger on top of the other (Bays and Wolpert, 2008; Kilteni and Ehrsson, 2017b, 2017a; Kilteni et al., 2018). This shows that only motor commands that reliably predict self-generated tactile stimuli produce robust somatosensory attenuation. Therefore, in our experiment, we also included distance between the hands as an additional experimental factor to further control for the mere effect of the simultaneous presence of movement and touch. This factor was not considered in the previous studies (Blakemore et al., 1998) but is valuable to control for it because it involves effects potentially related to splitting of attention (to both hands), sense of agency, and general cognitive anticipation of tactile stimulation.

We hypothesized that the attenuation of self-generated touch applied on the left index finger would be related to activity in the left cerebellum – that is, ipsilateral to the passive limb – compared to the control conditions. Moreover, we predicted that the degree of functional connectivity between the cerebellum and somatosensory areas would predict the degree of behaviorally estimated somatosensory attenuation across participants. Our results provide support for both of these hypotheses, which collectively provide strong evidence that the cerebellum plays a critical role in the attenuation of self-generated touch through its connectivity with somatosensory cortical areas.

## Materials and Methods

### Participants

After providing written informed consent, thirty naive participants (15 women and 15 men, 28 right-handed and 2 ambidextrous) aged 20-39 years participated in the study. Handedness was assessed using the Edinburgh Handedness Inventory (Oldfield, 1971). The sample size was set based on previous studies (Blakemore et al., 1998; Shergill et al., 2013) after taking into account the increased number of conditions in the present study. The Regional Ethical Review Board of Stockholm approved the study. After preprocessing of the fMRI scans, two participants were excluded due to motion artifacts. To be consistent, these two participants were also excluded from the behavioral study. Therefore, both behavioral and fMRI analyses were performed with a total of 28 participants.

### Procedures for the psychophysics task

The psychophysics task was performed approximately 30 minutes after the end of the fMRI experiment; this was the time it took to walk with the participants back from the scanner (Karolinska Hospital) to the psychophysics lab (Karolinska Institute). In the behavioral session, participants performed the classic force-matching task (Shergill et al., 2003). In each trial, the participants first received a force on the pulp of their left index finger by a probe controlled by a a DC motor (Maxon Motor EC 90 flat, manufactured in Switzerland) (*presented force*). A force sensor (FSG15N1A, Honeywell Inc., USA; diameter, 5 mm; minimum resolution, 0.01 N; response time, 1 ms; measurement range, 0–15 N) was placed inside the probe to measure the forces. After the application of each presented force, the participants used their right hand or index finger to produce a force on the left index finger (*matched force*) that matched the perceived intensity of the previously presented force. In two of the conditions (*press*_*0cm*_, *press*_*25cm*_), the participants reproduced the presented force by pressing their right index finger against a force sensor that was placed either on top of (but not in contact with) the probe (0 cm horizontal distance between the index fingers) or at a 25 cm distance from the probe (**Figure 1A** and **B**). This sensor controlled the force output of the lever with an approximately 30 ms intrinsic delay. In the third *slider* condition, the participants moved the wiper of a 13 cm slide potentiometer with their right hand (**Figure 1C**). As with the sensor, the slider controlled the force output on the participants’ fingers. The slider was positioned so that its midline laid at 25 cm to the right of the participants’ left index fingers. The lower limit of the slider (left extreme) corresponded to 0 N and the maximum (right extreme) to 5 N. Each trial started with the slider at 0 N. This *slider* condition is a classical control condition known to not involve somatosensory attenuation but used to assess basic somatosensory perception.

**Figure 1.**
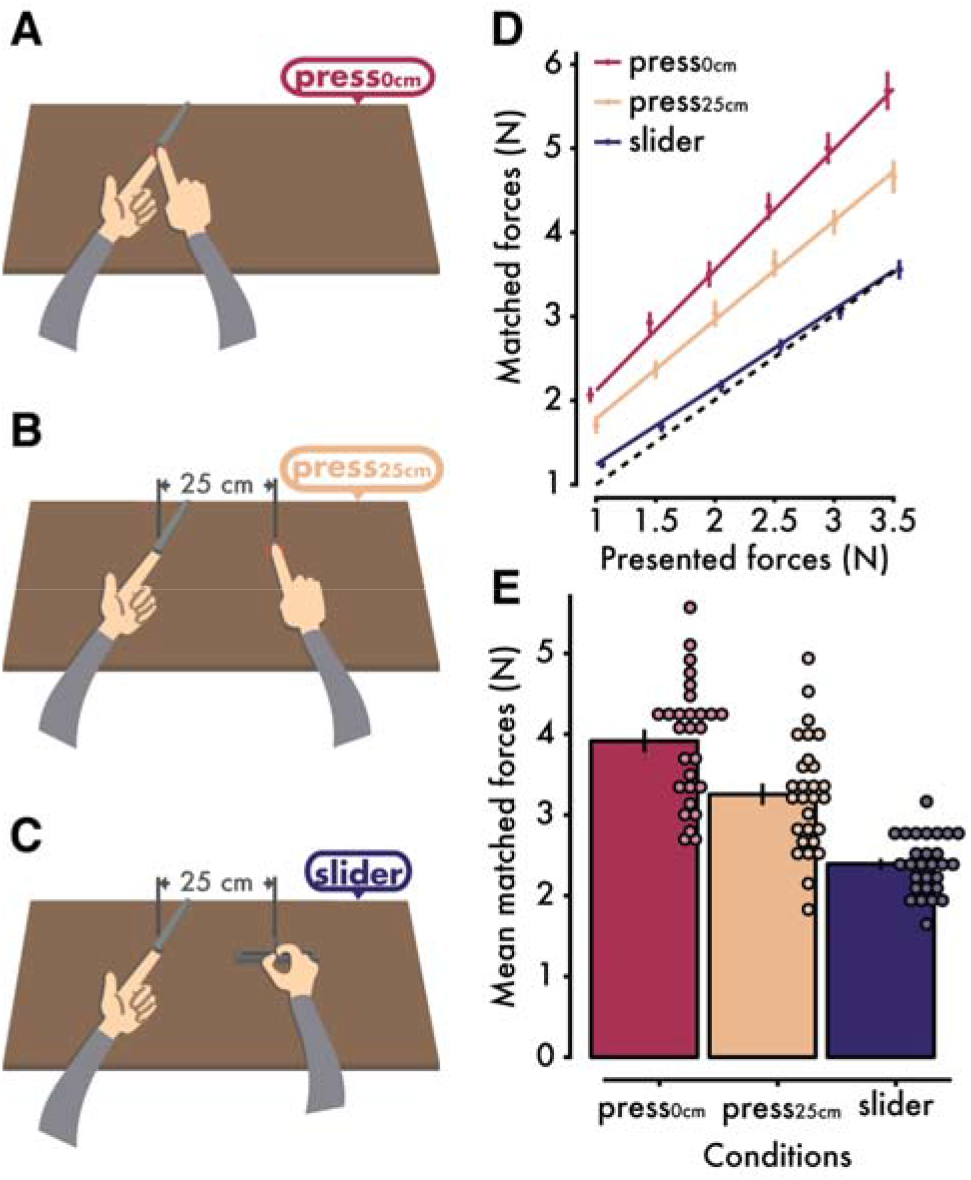
Experimental conditions and psychophysics results quantifying somatosensory attenuation. In each trial, participants received a force on their left index finger by a probe attached to a lever controlled by a motor. A force sensor inside the probe measured the applied force. Immediately afterwards, the participants had to reproduce the same force by pressing their right index finger against a sensor that controlled the force output on their left index finger. The sensor was placed either (**A**) on top of their left index finger (*press*_*0cm*_ condition) or (**B**) at 25 cm to the right of their left index finger (*press*_*25cm*_ condition). In the slider condition (**C**), participants reproduced the force by moving with their right hand a slider that controlled the force output on their left index finger. (**D**) Forces generated by the participants (*matched forces*) as a function of the externally generated forces (*presented forces*) (mean ± *SE* across participants). The dotted line represents theoretically perfect performance, and the colored lines are the fitted regression lines for each condition. The position of the markers has been horizontally jittered for visualization purposes. (**e**) Mean matched forces (± *SE*) per condition. The matched forces were significantly stronger in the *press*_*0cm*_ condition than in the other two conditions, meaning that the strongest attenuation of self-generated touch occurred when the hands simulated direct contact (i.e., no lateral distance) (**Figure 1**–**1**). Individual data points are overlaid onto the bars per condition.

Each of the three experimental conditions (*press*_*0cm*_, *press*_*25cm*_ and *slider*) consisted of 36 trials, with each level of the presented force (1 N, 1.5 N, 2 N, 2.5 N, 3 N and 3.5 N) pseudorandomly presented six times. To control for any order effects, the order of the conditions was fully counterbalanced across participants. During all conditions, the participants wore headphones through which white noise was presented to preclude the possibility that any noise generated by the motor served as a cue for the task. Auditory ‘go’ and ‘stop’ signals notified participants when to start and stop reproducing the presented force. A mark on the wall served as the participants’ fixation point. The forces applied by the motor (*presented force*) lasted 3 seconds, and participants had 3 seconds to reproduce the perceived force (*matched force*). The next force was presented approximately 3 seconds after the end of the previous matched force. No feedback was provided to the participants concerning their performance.

### Processing and statistical analysis of psychophysical data

We calculated the average of the matched force data produced on the left index finger at 2000–2500 ms after the ‘go’ signal to ensure that the force level had stabilized and the participants had not yet started to release the sensor (Bays and Wolpert, 2008; Kilteni and Ehrsson, 2017a, 2017b). The matched forces were then averaged across the six repetitions of each force level presented.

Two trials (out of 36) corresponding to two repetitions of two different force levels were missing for one participant in one experimental condition. For two different participants, one repetition of one force level was missing, and another was accidentally repeated in one of the three experimental conditions.

The psychophysics data were processed with Python (version 2.7.10) and analyzed using R (version 3.5.3). A repeated-measures analysis of variance (ANOVA) with the presented force level (1 N, 1.5 N, 2 N, 2.5 N, 3 N, 3.5 N) and the condition (*press*_*0cm*_, *press*_*25cm*_ and *slider*) as factors was used to analyze the matched forces. Planned pairwise comparisons were performed using either paired t-tests or paired Wilcoxon signed-rank tests, depending on whether the data satisfied normality assumptions.

### fMRI data acquisition

fMRI acquisition was performed using a General Electric 3T scanner equipped with an 8-channel head coil. T2-weighted echo-planar images (EPIs) containing 42 slices were acquired (repetition time: 2000 ms; echo time: 30 ms; flip angle: 80°; slice thickness: 3 mm; slice spacing: 3.5 mm; matrix size: 76 × 76; in-plane voxel resolution: 3 mm). A total of 1460 functional volumes were collected for each participant (365 volumes per run). For the anatomical localization of activations, a high-resolution structural image containing 180 slices was acquired for each participant before the acquisition of the functional volumes (repetition time: 6404 ms; echo time: 2.808 ms; flip angle: 12°; slice thickness: 1 mm; slice spacing: 1 mm; matrix size: 256 × 256; voxel size: 1 mm × 1 mm × 1 mm).

### Procedures for the fMRI experiment

The fMRI experiment always proceeded the force-matching task to keep participants blind to the experimental hypotheses. During the MRI session, participants laid comfortably in a supine position on the MRI scanner bed with their left hands placed palm-up on an MR-compatible plastic table (**Figure 2A**). Their left index finger was in contact with a 3D-printed probe that contained a force sensor (same specifications as above) and that was controlled by a motor (Maxon DC Motor RE40; reference 148866; manufactured in Switzerland) through string-based transmission. The string was tensioned through a pulley system consisting of ceramic bearings, and the transmission was mounted over a wooden structure with 6 degrees of freedom. Participants had their right index finger next to a second force sensor that was also placed on the table, either on top of (but not in contact with) the probe on the left index finger or at a 25 cm distance from it (**Figure 2A**). Sponges were used to support the participants’ arms in a comfortable posture inside the scanner so that they could keep their hands and fingers relaxed. Participants were instructed to fixate on the fixation cross displayed though a mirror screen that was mounted on the head coil (**Figure 2B**).

**Figure 2.**
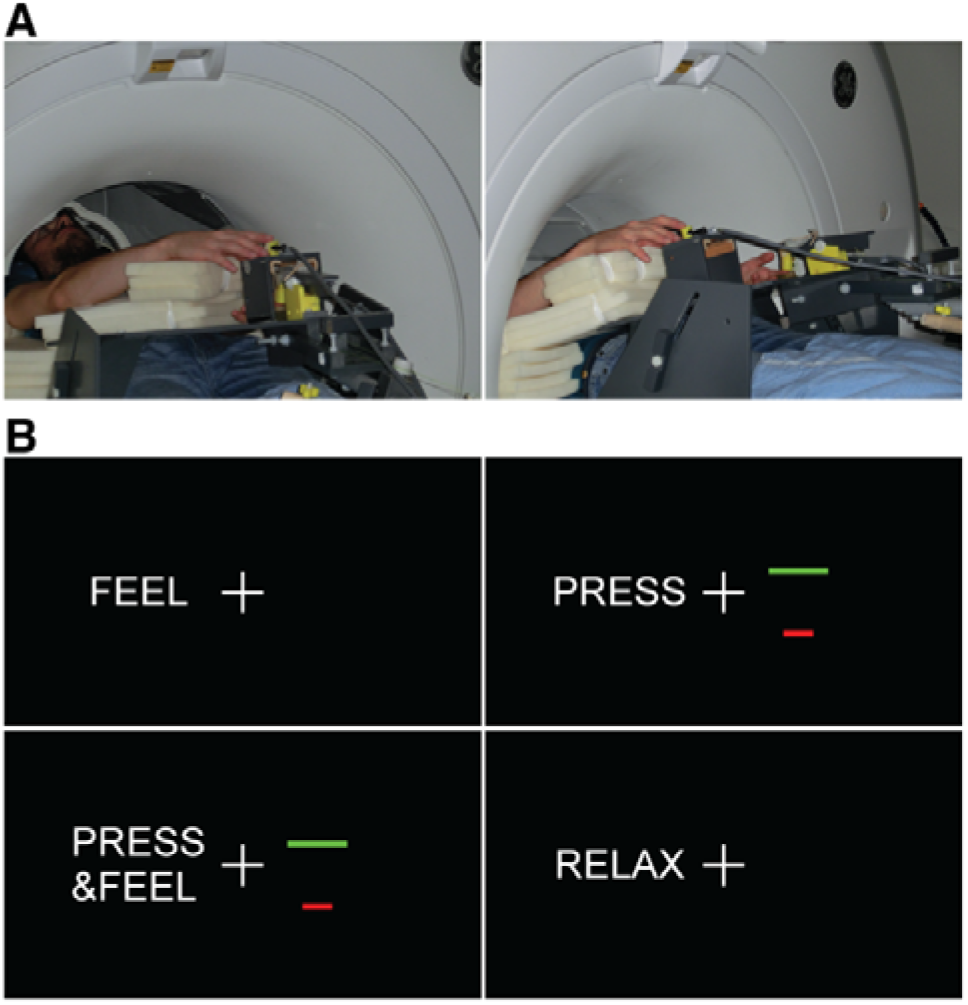
FMRI Experimental setup and instructions. **(A)** In two of the runs, the participants had their hands vertically aligned without any horizontal distance (0 cm), simulating direct contact (left), while in the remaining two runs, the participants’ hands were horizontally displaced by 25 cm (right). **(B)** The messages that participants received on the screen indicated the different conditions. See also **Figure 2-1**.

The DC motor controlling the lever was shielded inside a custom-made box made of *mu* metal and placed within a larger aluminum box. The motor box was placed inside the MRI room as far as possible from the scanner, and it was screwed to the hospital furniture for safety reasons. The motor cable was fitted with ferrite sleeves and passed through a hole to the control room where it was powered. Signal-to-Fluctuation-Noise Ratio tests ensured that the presence of the motor in the room did not produce any degradation in the quality of the MR images.

We used a factorial block design with the following three within-subjects’ factors: the *movement* of the right index finger versus no movement, the *touch* on the left index finger versus no touch, and the *distance* between the hands being either 0 cm or 25 cm. The design resulted in eight conditions: self-generated touch_0cm_, self-generated movement_0cm_, external touch_0cm_, rest_0cm_, self-generated touch_25cm_, self-generated movement_25cm_, external touch_25cm_ and rest_25cm_ (**Table 1**) (see below for an explanation of the task associated with each condition).

**Table 1.** Experimental factors and conditions in the fMRI experiment.

|  | No distance (0 cm) |  | Distance (25 cm) |  |
| --- | --- | --- | --- | --- |
|  | Touch | No touch | Touch | No touch |

| **Self-generated movement** | *self-generated touch<sub>0cm</sub>* | *self-generated movement<sub>0cm</sub>* | *self-generated touch<sub>25cm</sub>* | *self-generated movement<sub>25cm</sub>* |
| --- | --- | --- | --- | --- |
| **No self-generated movement** | *external touch<sub>0cm</sub>* | *rest<sub>0cm</sub>* | *external touch<sub>25cm</sub>* | *rest<sub>25cm</sub>* |

There were 4 runs: two were performed with the participants’ hands at a horizontal distance of 0 cm and two with a 25 cm distance introduced. Within each run, the participants performed the conditions *self-generated touch*, *self-generated movement*, *external touch* and *rest* at the corresponding distance. Each condition lasted 15 seconds. A 15-second rest period between conditions allowed the BOLD signal to return to baseline. These rest periods were not modeled in the analysis but served as an implicit baseline. Each of the four conditions was repeated 6 times within each run, resulting in a 12-minute run. The order of conditions was randomized both within and between participants. The order of the runs with respect to the distance factor was fully counterbalanced across participants.

Participants received visual instructions about the task in each condition on a screen seen via a mirror (**Figure 2B**). The message ‘*feel*’ indicated an externally applied force (2 N) on their left index finger (conditions: *external touch*). The message ‘*press’* instructed participants to press the sensor with their right index finger, as strongly as needed to increase the height of a red bar and make it reach a green line limit corresponding to 2 N (conditions: *self-generated movement*); no touch was felt on the left index finger in these conditions. The message ‘*press&feel*’ prompted participants to press the sensor with their right index finger (2 N) so that the red bar reached the green line, but in this condition, the participants simultaneously felt their self-generated touch on their left index finger (conditions: *self-generated touch*). Finally, the message ‘*relax’* asked participants to relax their hands (conditions: *rest*).

### Preprocessing of fMRI data

Functional data were preprocessed using the CONN toolbox (version 18a) (Whitfield-Gabrieli and Nieto-Castanon 2012) in SPM 12. Images were realigned, unwarped and slice-time corrected. Outlier volumes were detected using the Artifact Detection Tools employing the option for liberal thresholds (global-signal threshold of *Z* = 9 and subject-motion threshold of 2 mm). Then, the images were simultaneously segmented into gray matter, white matter and cerebrospinal fluid and normalized into standard MNI space (Montreal Neurological Institute, Canada). As a final step, the images were spatially smoothed using an 8 mm FWHM Gaussian kernel. The structural images were also simultaneously segmented (into gray and white matter and cerebrospinal fluid) and normalized to MNI space.

For the functional connectivity analysis, data were further denoised using the component-based noise correction method (*CompCor*) as it is implemented in the CONN toolbox. Five principal components from white matter, five principal components from cerebrospinal fluid, twelve principal realignment components (six plus 1^st^ order derivatives) and scrubbing parameters, together with two principal components per condition (the time series and its first derivative), were extracted and used as confounds. A bandpass filter [0.008, 0.09 Hz] was applied, and the data were linearly detrended.

### Statistical analysis of fMRI activations

After preprocessing, the data were analyzed with a general linear model (GLM) for each participant in Statistical Parametric Mapping 12 (SPM12; Welcome Department of Cognitive Neurology, London, UK, http://www.fil.ion.ucl.ac.uk/spm). Regressors were included for each of the eight conditions in the four scanning runs. In addition, the six motion parameters and any outlier volumes were included as regressors of no interest. Each condition was modeled with a boxcar function and convolved with the canonical hemodynamic response function of SPM 12. Contrasts of each condition regressor against zero were created.

At the second level of analysis, random-effects group analyses were performed by entering the contrast images of the condition regressors from each subject into two complementary full-factorial models. The first factorial model tested for the attenuation of self-generated touch compared to externally generated touch. For this model, we used the four condition regressors that corresponded to a distance of 0 cm (self-generated touch_0cm_, self-generated movement_0cm_, external touch_0cm_, rest_0cm_), and we inserted two repeated factors with unequal variance: the *movement* of the right index finger and the *touch* on the left index finger. A second factorial model was created to assess the effect of distance on the attenuation of self-generated touch. For this model, we used the condition regressors of all movement conditions (self-generated touch_0cm_, self-generated movement_0cm_, self-generated touch_25cm_, self-generated movement_25cm_), and we inserted two repeated factors with unequal variance: the *touch* on the left index finger and the *distance* between the hands (0 cm or 25 cm).

Contrasts of interest focused on the interaction effects of each factorial model. Specifically, the *Movement*_*0cm*_ -*by*- *Touch*_*0cm*_ interaction effect, i.e., (self-generated movement_0cm_ + external touch_0cm_ − self-generated touch_0cm_ − rest_0cm_) > 0, was calculated to investigate the attenuation of self-generated touch compared to externally generated touch after factoring out the main effects of movement and touch. This contrast allows to study the attenuation of touch on the passive left index finger that was produced by the moving right index finger, but importantly without the concomitant effects of the movement of the right hand. Similarly, the *Touch -by-Distance* interaction, i.e., (self-generated touch_25cm_ + self-generated movement_0cm_ − self-generated touch_0cm_ − self-generated movement_25cm_) > 0, served to distinguish the attenuation of self-generated touch from a condition that involved the simultaneous presence of movement and touch but no robust somatosensory predictions. Both directions of interaction effects as well as all the main effects are reported for clarity and transparency.

Given our strong a priori hypotheses about cerebellar and somatosensory areas in the corresponding 2-by-2 interactions, we applied a correction for multiple comparisons in all statistical tests within such regions of interest. Specifically, two cerebellar regions of interest were defined as spheres centered around the cerebellar peak found in the study of Blakemore et al. (1998) [MNI coordinates: x = 22, y = −58, z = −22] and its ipsilateral analogue, i.e., that derived by flipping the x-coordinate [x = −22, y = −58, z = −22]. These coordinates were originally specified in MNI space [Sarah-Jayne Blakemore, personal communication, November 28, 2018], and therefore, they were not converted from Talairach space. Somatosensory regions of interest were defined as spheres centered around the corrected or uncorrected primary and secondary somatosensory cortical peaks detected from the main effects of touch. Since our factorial designs were balanced, the main effects and interactions are orthogonal contrasts, ensuring no statistical inference bias and allowing us to use the main effects as functional localizers (Friston et al., 2006). Given the somatotopic specificity of somatosensory areas, we used spheres of 10-mm radius for defining somatosensory regions of interest. In contrast, given earlier findings assigning sensorimotor hand-related functions to several cerebellar areas, including lobules V, VI and Crus I (Blakemore et al., 1998, 2001; Diedrichsen et al., 2005; King et al., 2018), the two cerebellar spheres were set to have a 15-mm radius to include a larger cerebellar volume. Statistical tests for main effects were corrected for multiple comparisons over the entire brain.

For each peak activation, the coordinates in MNI space, the *z* value and the *p* value are reported. We denote that a peak survived a threshold of *p* < 0.05 after correction for multiple comparisons at the whole-brain or small volume by the term “FWE-corrected” following the *p* value. Alternatively, the term “uncorrected” follows the *p* value in the few cases when the activation did not survive correction for multiple comparisons, but it is still informative to describe. For example, cerebellar peaks that are outside the regions of interest and did not survive corrections for multiple comparisons are still informative to report for descriptive purposes. However, all main results on which our main conclusions are drawn survived corrections for multiple comparisons.

### Anatomical labeling and visualization of the results

We only reported peaks of clusters that had a size larger than 3 voxels and were situated within gray matter. For labeling the anatomical localizations of the significant peaks of activation, we used the nomenclature from the human brain atlas of Duvernoy (1999). For labeling the anatomical localization of cerebellar peaks of activation, we used the probabilistic atlas of the cerebellum provided with the SUIT toolbox (Diedrichsen et al. 2009) and included in the Anatomy toolbox (Eickhoff et al. 2005) after specifying that the normalization be performed using the MNI template; we labeled the peaks according to the area for which they showed the highest probability. If the probability given for the cerebellar area was within 40-60%, we also reported the area that showed the next highest probability. Activations driven by main effects were rendered on the standard single subject 3D-volume provided with SPM for an overview of the activation pattern in the whole brain. Peaks from both main and interaction effects that were important for our hypotheses were overlaid onto the average anatomical image for all participants in the study to facilitate precise anatomical localization. For better visualization of the cerebellar peaks, the thresholded maps of the cerebellar activations were overlaid onto the cerebellar flatmap (glass-brain projection) provided by the SUIT toolbox, after specifying that volume-based normalization was done in SPM (Diedrichsen and Zotow 2015). To isolate the cerebellar peaks from the rest of the brain when needed, we applied an anatomical mask over the entire cerebellum (both vermis and hemispheres) that was created with the Anatomy toolbox. For visualization purposes and to access the anatomical specificity of our effects in a purely descriptive manner, all activation maps are displayed at a threshold of *p* < 0.001 uncorrected.

### Statistical analysis of fMRI connectivity

A seed-to-voxel analysis was conducted in the form of generalized psychophysiological interactions (McLaren et al., 2012) using the denoised data within the CONN toolbox. Seeds were defined as spheres with a 10-mm radius around the cerebellar and somatosensory peaks revealed by the activation analysis (interaction contrasts). At the group level, the contrasts of interest consisted of the *Movement*_*0cm*_ -*by*- *Touch*_*0cm*_ interaction effect – i.e., (self-generated touch_0cm_ + rest_0cm_ − self-generated movement_0cm_ − external touch_0cm_) > 0 – that assesses connectivity increases during the self-generated touch condition compared to external touch after factoring out the main effects, and the *Touch -by- Distance* interaction – i.e., (self-generated touch_0cm_ + self-generated movement_25cm_ − self-generated touch_25cm_ − self-generated movement_0cm_) > 0 – that assesses connectivity increases during the self-generated touch condition compared to the simultaneous presence of movement and touch after factoring out the main effects. Since we were interested in the attenuation of self-generated touch, we only assessed increases – and not decreases – in brain connectivity in the self-generated condition compared to control conditions.

To identify connectivity changes that were specific to somatosensory attenuation, we used as a second-level covariate the participants’ attenuation index as extracted from the force-matching task. For each participant we calculated the difference between the mean force he/she exerted in the condition of interest and the force that he/she exerted in a reference condition, similar to our previous study (Kilteni and Ehrsson, 2017b). Specifically, to investigate connectivity increases in the self-generated touch condition compared to the externally generated touch condition (*Movement*_*0cm*_ -*by*- *Touch*_*0cm*_ interaction), we used the difference between the mean matched force in the *press*_*0cm*_ condition and the mean matched force in the *slider* condition. Analogously, to investigate connectivity increases in the self-generated touch condition with respect to the simultaneous movement and touch condition (*Touch -by- Distance* interaction), we used the difference between the mean matched force in the *press*_*25cm*_ condition and that in the *press*_*0cm*_ condition. By doing so, the contrasts of brain activation were ‘aligned’ with the behavioral contrasts, allowing a proper covariate analysis. We tested for connectivity changes both between somatosensory and cerebellar areas as well as between different areas within the cerebellum. Statistical maps were assessed using corrections for multiple comparisons, as described above.

## Results

### Behavioral attenuation of self-generated forces

**Figure 1D** shows the participants’ performance per condition and presented force level. A repeated-measures ANOVA revealed a significant main effect of condition (*F*(2, 54)□=□121, *p*□<□0.001, *η*^2^ = 0.020), a significant main effect of the presented force level (*F*(5, 135)□= 414.3□, *p*□*>*□0.001, *η*^2^ = 0.521), and a significant interaction (*F*(10, 270)□=□15.23, *p*□*<*□0.001, *η*^2^ = 0.017). Pairwise comparisons between the levels of the presented forces revealed significant differences for each pair (all *p*-values□*<*□0.001), confirming that the participants clearly discriminated each presented force level.

Importantly, as seen in **Figure 1D** and **E**, the participants produced stronger forces when their hands were horizontally aligned (mean□±□SD□=□3.915□±□0.752□N) than when they were spatially separated (mean□±□SD□=□3.255□±□0.711□N) or when they used the slider to reproduce the forces (mean□±□SD□=□2.392□±□0.357□N). Pairwise comparisons revealed significant differences between the *press*_*0cm*_ and the *press*_*25cm*_ conditions (*t*(27)□=□8.63, *p*□*<*□0.001, 95% confidence interval *CI*□=□[0.503, □0.817], Cohen’s *d* = 1.631), between the *press*_*0cm*_ and the *slider* conditions (*t*(27)□=□13.57, *p*□*<*□0.001, *CI* □=□[1.293, □1.754], Cohen’s *d* = 2.564) and between the *press*_*25cm*_ and the *slider* conditions (*t*(27)□=□8.43, *p*□*<*□0.001, *CI* □=□[0.65, □1.07], Cohen’s *d* = 1.593) (**Figure 1-1)**. Taken together, these findings replicate previous results indicating strong attenuation when the hands simulate direct contact and significantly reduced attenuation when the hands are spatially separated or when a slider is used to reproduce the force (Bays and Wolpert, 2008; Kilteni and Ehrsson, 2017a, 2017b; Kilteni et al., 2018).

### Behavioral performance inside the scanner

It is important to confirm that the participants performed the fMRI tasks as requested; that is, that they applied and received the required intensity of forces (2 N). By confirming this, we can ensure that any differences in the BOLD signals were not due to different levels of force being experienced in the different conditions (Ehrsson et al. 2001). To this end, we analyzed the data from the left and the right index finger sensors collected from the fMRI sessions. We considered only the last 10 seconds (and not the entire 15 seconds) of each condition to account for the participants’ reaction time to press the sensor and for the initial period when they were adjusting the force before reaching the desired level of the target force.

With respect to the left index finger sensor, a repeated-measures analysis of variance (ANOVA) with the factors of *distance* (0 cm or 25 cm) and *mode* (self-generated or externally generated) revealed no significant effect of distance (*F*(1, 27)□=□0.06, *p*□*=*□0.808, *η*^2^ < 0.001), no effect of mode (*F*(1, 27)□=□0.47, *p*□*=*□0.499, 2 *η*^2^ = 0.007) and no significant interaction between them (*F*(1, 27)□=□0.05, *p*□*=*□0.820, *η*^2^ < 0.001). A Bayesian repeated-measures ANOVA using JASP (2019) revealed that the data were 61.95 times more likely to occur under the null model (i.e., a model not including the effects of distance, mode and their interaction) compared to a model including these effects (**Figure 2-1**).

With respect to the right index finger sensor, a repeated-measures ANOVA with the factors of *distance* (0 cm or 25 cm) and *mode* (self-generated movement or self-generated touch) revealed no significant effect of distance (*F*(1, 27)□=□0.10, *p*□*=*□0.758, *η*^2^ < 0.001), no effect of mode (*F*(1, 27)□=□0.21, *p*□*=*□0.653, *η*^2^ < 0.001) and no significant interaction between these factors (*F*(1, 27)□=□0.02, *p*□*=*□0.880, *η*^2^ < 0.001). A Bayesian repeated-measures ANOVA revealed that the data were 81.98 times more likely to occur under the null model (i.e., a model not including the effects of distance, mode and their interaction) compared to a model including these effects (**Figure 2-1**).

In conclusion, the above analysis eliminated the possibility that any force differences could account for our fMRI findings – a factor that was not controlled in earlier studies on somatosensory attenuation (Blakemore et al., 1998, 2001).

### Neural attenuation of self-generated touch compared to externally generated touch

We first tested for the attenuation of self-generated touch compared to externally generated touch by building a 2-by-2 factorial model that included the four experimental conditions that corresponded to the distance of 0 cm (see *Materials and Methods*). The interaction term of such a model represents the difference between externally generated and self-generated touch, critically after factoring out activity that is due to the main effect of movement or the main effect of touch alone. This design further allowed a direct comparison between our data and the data of Blakemore et al. (1998).

As expected, the main effect of moving the right index finger revealed widespread activity in several areas, including the left primary motor cortex (M1), dorsal (PMd) and ventral (PMv) premotor cortex, supplementary motor area, and putamen and the right cerebellum (**Figure 3-1**, **Table 2-1**). The main effect of tactile stimulation on the left index finger was associated with activations in the right parietal operculum (putative secondary somatosensory cortex, S2) and the right and left supramarginal gyri (SMG) in the inferior parietal lobule (**Figure 3-2**). Situated in the inferior parietal lobe, the supramarginal gyrus is part of the sensory association cortex and is involved in higher-order somatosensory processing (Bodegård et al., 2001; Lamp et al., 2019). At the uncorrected level of *p* < 0.001, the right primary somatosensory cortex (S1) was also activated (**Table 2-2**).

**Figure 3.**
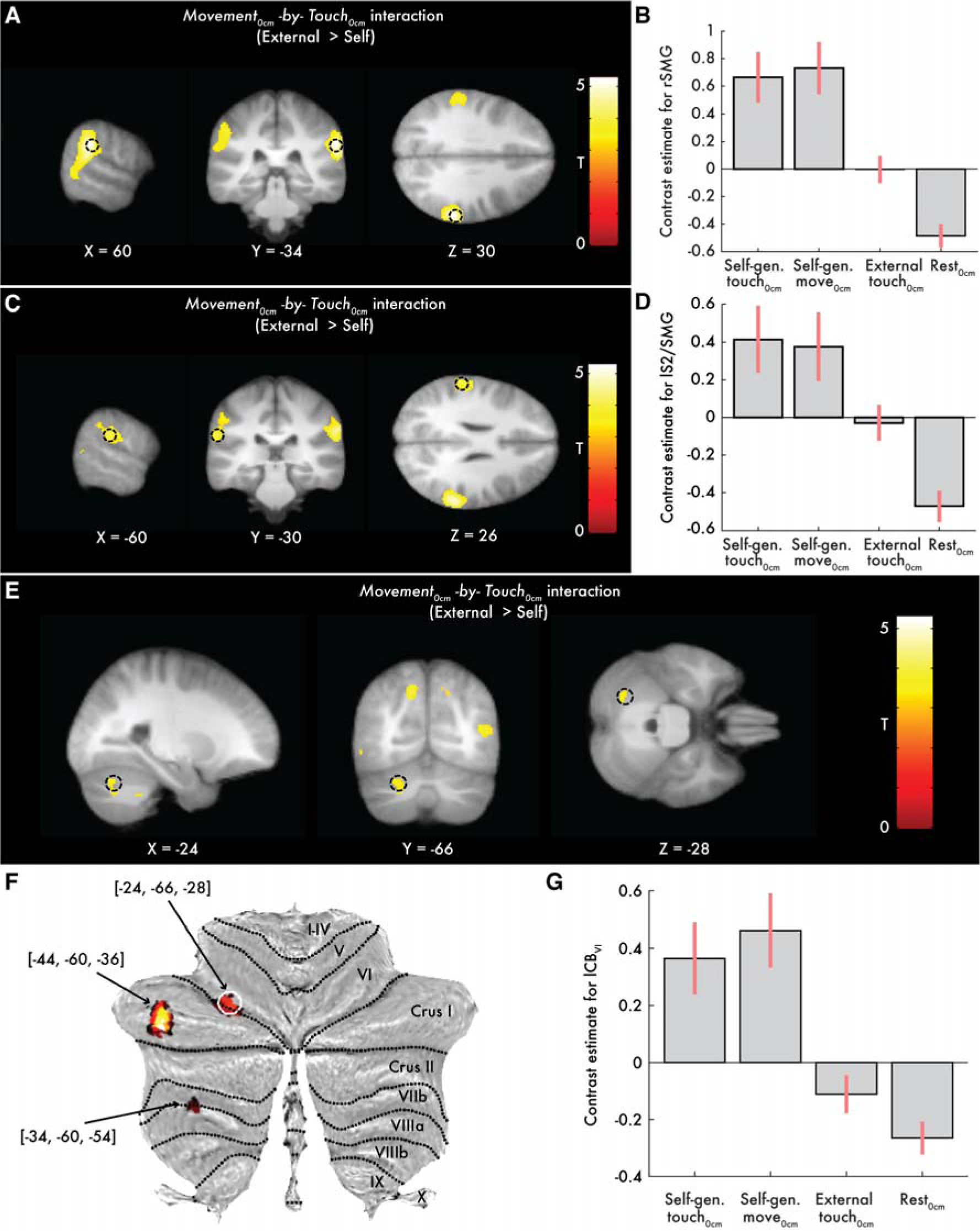
Somatosensory and cerebellar activations revealed by the *Movement*_*0cm*_ -*by*- *Touch*_*0cm*_ interaction. (Direction: External > Self). Activations reflect greater effects when the touch is delivered in the absence of movement (BOLD change from the *rest*_*0cm*_ to the *external touch*_*0cm*_ condition) than during a self-generated movement (BOLD change from the *self-generated movement*_*0cm*_ to the *self-generated touch*_*0cm*_ condition). **(A**, **C**, **E)** Slice views of significant peaks (*p* < 0.05 FWE-corrected) at the right and left supramarginal gyri (next to S2) and left cerebellum, indicated by black circles. The activations (here and in all subsequent figures unless stated otherwise) have been overlaid on the average anatomical image of the participants. (**F**) Cerebellar activations overlaid onto a cerebellar flatmap. The peak in lobule VI indicated by the white circle survived FWE corrections. For descriptive purposes, two more peaks of left posterior cerebellar clusters are also shown at the uncorrected level of *p* < 0.001 (**Table 2-3**). The largest activation was observed in the middle of lobule Crus I but that posterior activation did not survive corrections for multiple comparisons. No activation peaks were detected in the right cerebellar hemisphere, not even at the threshold of *p* < 0.001 uncorrected. **(B**, **D**, **G)** Bar plots of the contrast estimates per condition and peak in arbitrary units. Error bars denote 90% confidence intervals. All activation maps were thresholded at *p* < 0.001 uncorrected for visualization purposes and to descriptively illustrate the anatomical specificity of the significant effects. See also **Figures 3-1, 3-2** and **3-3**.

**Table 2.** Activation peaks for the *Movement*_*0cm*_ -*by*- *Touch*_*0cm*_ interaction. Peaks reflecting greater effects during touch in the absence of movement compared to touch in the context of a self-generated movement (Direction: External > Self). See also **Tables 2-1, 2-2, 2-3, 2-4, 2-5** and **2-6**.

| Brain region | Cluster size (voxels) | MNI coordinates (mm) |  |  | z | p |
| --- | --- | --- | --- | --- | --- | --- |
|  |  | x | y | z |  |  |
| R supramarginal gyrus | 1637 | 60 | -34 | 30 | 4.97 | $p = 0.007$ FWE-corrected |
| R temporal parietal junction | | 62 | -36 | 22 | 4.64 | $p = 0.027$ FWE-corrected |
| L parietal operculum (S2) /supramarginal gyrus | 221 | -60 | -30 | 26 | 3.92 | $p = 0.002$ FWE-corrected* |
| L supramarginal gyrus | | -58 | -22 | 26 | 3.88 | $p = 0.003$ FWE-corrected* |
| L supramarginal gyrus | | -56 | -34 | 32 | 3.60 | $p = 0.007$ FWE-corrected* |
| L cerebellum VI | 44 | -24 | -66 | -28 | 3.47 | $p = 0.026$ FWE-corrected* |

\* After small volume correction

When testing the *Movement*_*0cm*_ -*by*- *Touch*_*0cm*_ interaction that reveals effects related to somatosensory attenuation, significant peaks (*p* < 0.05 FWE-corrected) were detected at the right supramarginal gyrus next to S2, the junction between the right superior temporal gyrus and supramarginal gyrus, the junction between the left parietal operculum (S2) and supramarginal gyrus, the left supramarginal gyrus and the left cerebellum (lobule VI) (**Table 2**, **Figure 3**); all showed greater activation when the touch was delivered in the absence of movement (i.e., BOLD change from the *rest* to the *external touch* condition) than in the presence of a self-generated movement (i.e., BOLD change from the *self-generated movement* to the *self-generated touch* condition) (**Figure 3B** and **D**). No significant peaks were detected in the right cerebellum, even at the uncorrected level of *p* < 0.001 (**Table 2-3**, **Table 2-4**, **Table 2-5**, **Figure 3-3**). When examining the interaction contrast in the opposite direction, there were no significant peaks reflecting greater effects of self than externally generated touch in the *Movement*_*0cm*_ -*by*- *Touch*_*0cm*_ interaction (**Table 2-6**).

To examine which regions were responsible for driving the suppression of activity in somatosensory areas when the touch was delivered in the context of movement (self-touch), we conducted a generalized psychophysical interaction analysis (gPPI) to look for voxels in the whole brain that increased their functional connectivity with the peak at the right supramarginal gyrus (**Table 2**) during self-generated touch compared to external touch (*Movement*_*0cm*_ -*by*- *Touch*_*0cm*_ interaction, Direction: Self > External). Moreover, to isolate those connectivity changes that were specific to the somatosensory attenuation, we included the participants’ behavioral attenuation as a covariate in the analysis, i.e., each participant’s difference between the matched forces in the *press*_*0cm*_ and the *slider* condition in the force-matching task. Importantly, we found that the more the participants attenuated their self-generated forces in the force-matching task, the more the right supramarginal gyrus increased its connectivity with the left cerebellum (*p* < 0.05 FWE-corrected) (**Figure 4A-B**, **Table 3-1**, **Figure 4-1**, see also **Table 3-2**). Notably, when we removed the behavioral covariate, no voxels were detected in the cerebellum at the *p* < 0.001 uncorrected threshold, suggesting that the participants’ attenuation index was critical for this increased cerebrocerebellar connectivity.

**Figure 4.**
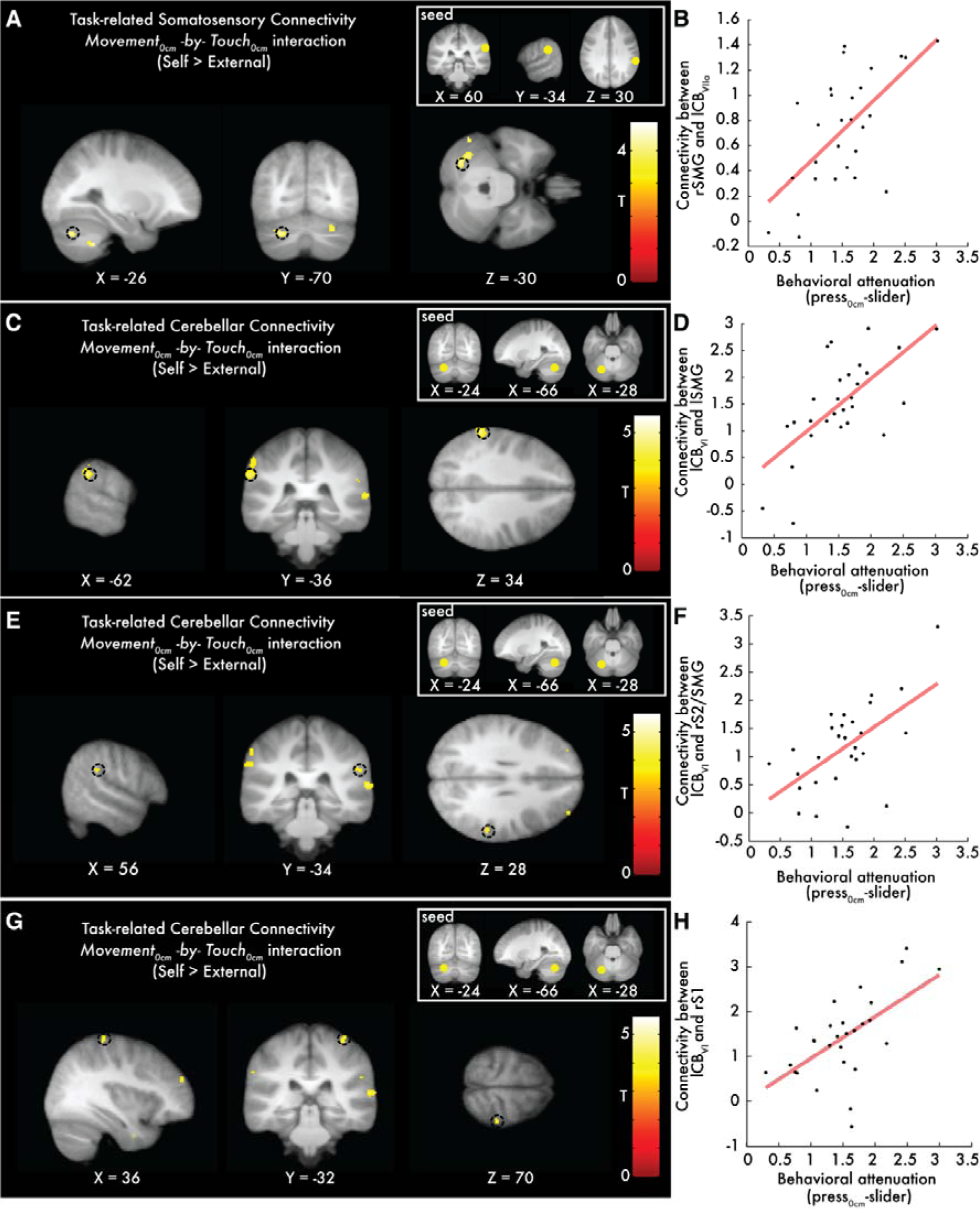
Cerebellar and somatosensory peaks showing increased connectivity with the seeds of interest as a function of behavioral attenuation. (**A**) Sagittal (left), coronal (middle) and axial (right) views of the significant peak in the left cerebellum (*p* < 0.05 FWE-corrected) that increased its connectivity with the right supramarginal gyrus (seed). Only the cerebellar peak denoted by the black circle was significant. (**B**) Scatterplot showing the relationship between the connectivity increases of the peak in (A) and the participants’ behavioral attenuation as measured in the force-matching task. (**C**, **E**, **G**) Slice views of the peaks in the left supramarginal gyrus, the right parietal operculum (S2)/supramarginal gyrus and the right primary somatosensory cortex (*p* < 0.05 FWE-corrected) that significantly increased their connectivity with the left cerebellum (seed, lobule VI), indicated by black circles. (**D**, **F**, **H**) Scatterplots showing the relationship between the connectivity increases of the peaks in (C, E, G) and the participants’ behavioral attenuation. See also **Figures 4-1** and **4-2**.

**Table 3.** Somatosensory cortical areas that increased their functional connectivity with the left cerebellum as a function of behavioral somatosensory attenuation. Peaks reflecting greater connectivity with the cerebellar seed during touch delivered in the context of a self-generated movement compared to touch delivered in the absence of movement as a function of behavioral attenuation (*Movement*_*0cm*_ -*by*- *Touch*_*0cm*_ interaction, Direction: Self > External). See also **Tables 3-1, 3-2, 3-3** and **3-4**.

| Brain region | Cluster size (voxels) | MNI coordinates (mm) |  |  | *z* | *p* |
| --- | --- | --- | --- | --- | --- | --- |
|  |  | x | y | z |  |  |
| **L supramarginal gyrus** | 26 | -62 | -36 | 34 | 3.88 | $p = 0.006$ FWE-corrected* |
| **R postcentral gyrus (S1)** | 14 | 36 | -32 | 70 | 3.37 | $p = 0.028$ FWE-corrected* |
| **R parietal operculum (S2)/ supramarginal gyrus** | 10 | 56 | -34 | 28 | 3.28 | $p = 0.036$ FWE-corrected* |

\* After small volume correction.

When the seed was placed in the left cerebellum, the gPPI analysis revealed increased cerebellar connectivity with both the left and right supramarginal gyri/parietal opercula (S2) and the right primary somatosensory cortex (S1) (*p* < 0.05 FWE-corrected), when the touch was self-generated compared to when it was externally generated (**Table 3**, **Figure 4C-H**, **Table 3-3**). We further observed connectivity increases to other regions within the cerebellum: bilateral peaks at lobules VII/VIII increased their connectivity with the seed at lobule VI the more the participants attenuated their self-generated forces in the force-matching task (**Figure 4-2**); these connectivity changes, however, did not survive corrections for multiple comparisons (*p* < 0.001 uncorrected threshold). When removing the participants’ individual behavioral attenuation as a covariate from the analysis, no significant increases (*p* < 0.001 uncorrected) were detected in the cerebellar connectivity with the somatosensory areas under discussion, and the intracerebellar effects disappeared (**Table 3-4**), which suggests that the functional connectivity under discussion is specifically related to somatosensory attenuation.

### Neural attenuation of self-generated touch compared to simultaneous movement and touch

The previous factorial design tested for the differential effects between self-generated and external touch, importantly after controlling for the main effects of movement and touch. However, it does not control for pure effects of bimanual actions involving the simultaneous presence of movement and touch, divided attention to the two hands and sense of agency, factors that could influence the BOLD signal in regions related to sensorimotor processing. Therefore, complementary to our previous analysis, we constructed a factorial design with the four self-generated conditions (self-generated touch_0cm_, self-generated movement_0cm_, self-generated touch_25cm_, self-generated movement_25cm_). This design controls for the effects described above and tests for neural attenuation of self-generated touch when the hands simulated direct contact (0 cm lateral distance) compared to when the hands were separated by 25 cm, leading to significantly reduced attenuation.

As expected, the main effect of tactile stimulation was associated with significant activation of the right parietal operculum (**Figure 5-1**) and, at the uncorrected level *p* < 0.001, the right primary somatosensory cortex (S1) (**Figure 5-2**). The main effect of distance revealed activity in motor-related areas, including the right and left precentral gyrus (M1) and the cerebellum (**Figure 5-3**, **Figure 5-4**, **Figure 5-5**), probably reflecting the difference in the postures of the arms in the distance manipulation (**Figure 1**).

**Figure 5.**
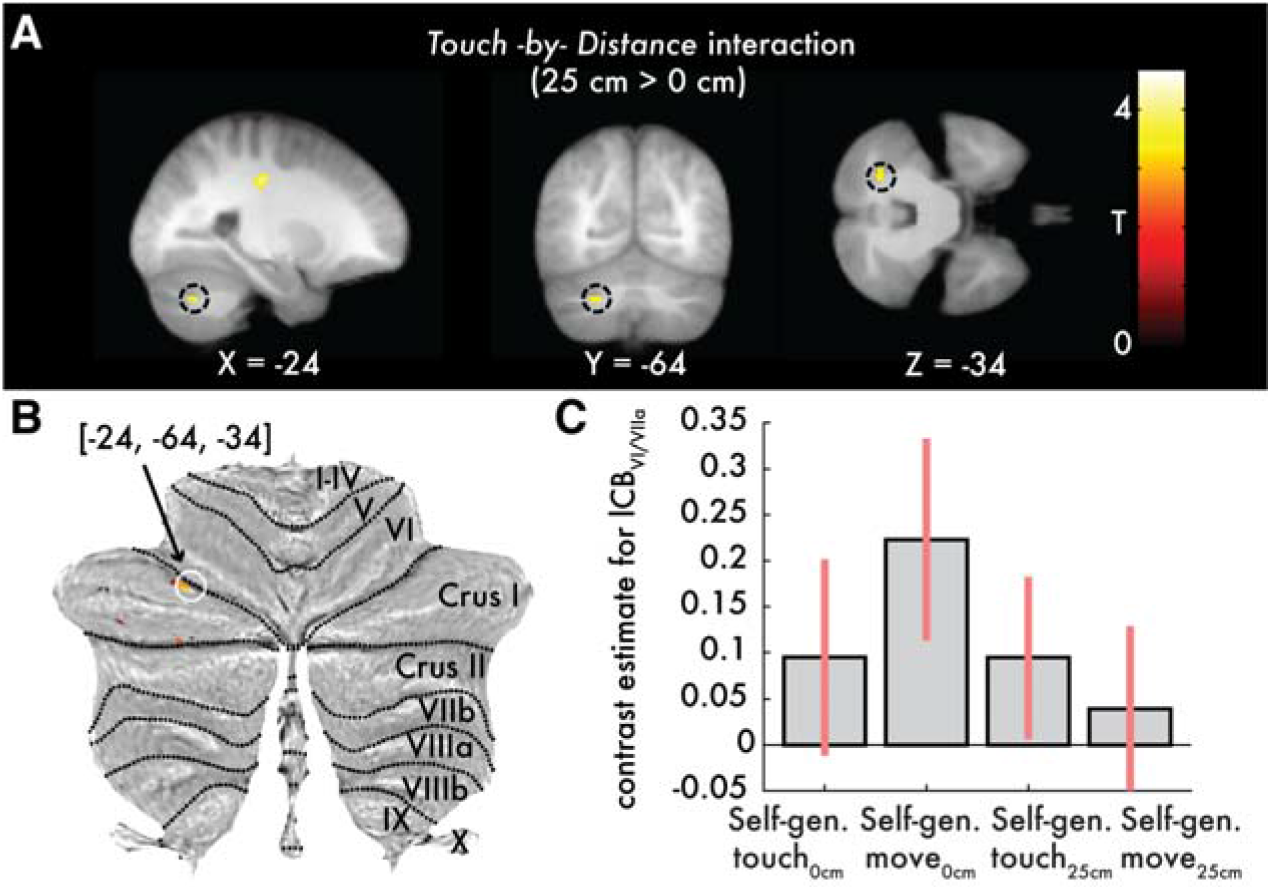
Cerebellar activations revealed by the *Touch -by- Distance* interaction. Activations reflecting greater BOLD responses when the touch is delivered in the presence of a 25 cm hands’ distance (BOLD change from the *self-generated movement*_*25cm*_ to the *self-generated touch*_*25cm*_ condition) than in the absence of distance between the hands (BOLD change from the *self-generated movement*_*0cm*_ to the *self-generated touch*_*0cm*_ condition) (Direction: 25 cm > 0 cm). (**A**) Slice views of the significant cerebellar peak (*p* < 0.05 FWE-corrected) denoted by the black circles in the sagittal (left), coronal (middle) and axial (right) planes, respectively. **(B)** Activations overlaid onto the cerebellar flatmap seen at the *p* < 0.001 uncorrected threshold. Only the peak denoted by the white circle survived FWE correction. There were no significant peaks in the right hemisphere at the *p* < 0.001 uncorrected level. (**C**) Bar plots of the contrast estimates per condition for the significant peak in arbitrary units. Error bars denote 90% confidence intervals. All activations are seen at *p* < 0.001 uncorrected for visualization purposes. See also **Figures 5-1, 5-2, 5-3, 5-4, 5-5, 5-6, 5-7, 5-8, 5-9, 5-10, 5-11** and **5-12**.

The important *Touch -by- Distance* interaction representing weaker activity when the self-generated touch is received with the hands being overlapping (0 cm distance) compared to when the hands are separated by 25 cm revealed significant effects in the left cerebellum (lobules VIIa Crus I/VI). Critically, the left cerebellum showed a suppression of activation in the absence of distance (i.e., BOLD change from the *self-generated movement*_*0cm*_ to the *self-generated touch*_*0cm*_ condition) than in the presence of distance (i.e., BOLD change from the *self-generated movement*_*25cm*_ to the *self-generated touch*_*25cm*_ condition) (*p* < 0.05 FWE-corrected) (**Figure 5, Figure 5-6**. No activations were detected in the right cerebellar hemisphere at the *p* < 0.001 uncorrected level (**Figure 5-7**, **Figure 5-8**). Moreover, no active voxels were observed for the *Touch -by- Distance* interaction in the opposite direction (self-generated touch_0cm_ > self-generated touch_25cm_) at the uncorrected level of *p* < 0.001.

Next, we looked for connectivity changes between the left cerebellar peak (seed, lobule VI/VIIa) and somatosensory areas using a whole brain gPPI analysis that included the participants’ attenuation index as a behavioral covariate, defined here as the difference between the matched forces in the *press*_*0cm*_ and *press*_*25cm*_ conditions. We found one peak of activation at the right postcentral gyrus (S1) that increased its connectivity with the cerebellum when the touch was presented in the absence of distance than in the presence of distance (*Touch -by- Distance* interaction, Direction: 0 cm > 25 cm) (**Figure 5-9**, **Figure 5-10)** as a function of the behaviorally registered attenuation across participants. Similarly, we observed that within the cerebellum, the more participants attenuated their self-generated forces, the stronger the connectivity between the seed at left lobule VI/VIIa and the anterior part of lobule VI bilaterally (**Figure 5-9**). However, none of these peaks survived corrections for multiple comparisons. Notably, when we removed the covariate, we no longer observed these cerebrocerebellar and intracerebellar effects (**Figure 5-11**).

Finally, we constructed the full factorial model with all three factors (Distance, Movement, Touch), and we calculated the three-way interaction using all eight conditions. This interaction reflects the difference between external and self-generated touch when the hands simulate direct contact (0 cm) compared to when the hands are apart (25 cm), after factoring out the three main effects and all the two-way interactions. Consistent with the results from our two two-way interaction analyses described above, this three-way interaction revealed significant activity in the left cerebellum (lobule VI) (*p* < 0.05 FWE-corrected) (**Figure 5-12**).

## Discussion

Using fMRI together with the classic force-matching task, we investigated the neural processes underlying the predictive attenuation of self-generated touch. We found that touch is associated with a suppression of activation in the bilateral secondary somatosensory cortex when presented in the context of a self-generated movement (self-generated touch) compared to touch of identical intensity that is presented in the absence of movement (externally generated touch), replicating previous results (Blakemore et al., 1998) and consistent with earlier findings on bilateral responses in these areas following unilateral stimulation (Eickhoff et al., 2008). In addition, we observed suppression of activation in the cerebellum during touch when presented in the context of a self-generated movement (self-generated) compared to the absence of movement and compared to a well-matched control condition involving the presence of distance between the hands. The site of this cerebellar activity was lateralized to the hemisphere that was *ipsilateral* to the passive limb that received the touch, i.e., the left, in contrast to the results of Blakemore et al. (1998) but in good agreement with the anatomical facts of an ipsilateral representation of the body in the cerebellum (Grodd et al., 2001; Manni and Petrosini, 2004) and the contralateral organization of the functional corticocerebellar pathways (O’Reilly et al., 2010; Buckner et al., 2011). Moreover, we found that functional connectivity between the ipsilateral cerebellum and the contralateral primary and bilateral secondary somatosensory areas increased during self-generated touch in a way that scaled linearly across participants with the somatosensory attenuation effect as quantified in the force-matching task. This observation is in contrast to that of Blakemore et al. (1999b), who reported functional correlations between the right cerebellum and right somatosensory areas that, given the cerebellar laterality, probably reflected processes related to the movement of the right hand rather than the somatosensory attenuation of the left hand. Together with other studies on sensory attenuation in the visual and/or auditory modalities (Knolle et al., 2013; Straube et al., 2017), our findings reveal the fundamental role of the cerebellum in predicting and cancelling self-generated somatosensory input. Moreover, they indicate that the functional connectivity between the cerebellum and the somatosensory cortex implement the somatosensory attenuation phenomenon.

What does this functional corticocerebellar coupling represent? By keeping in mind that functional connectivity between two areas does not necessarily imply a causal relationship (Eickhoff and Müller, 2015), one could hypothesize that this connectivity reflects the prediction signal that the cerebellum sends to somatosensory cortices to suppress their activity. Accordingly, given the copy of the motor command sent to the right hand, the cerebellum predicts contact of the right index finger with the left index finger, including the expected tactile feedback, and sends a cancelation signal to somatosensory areas to attenuate its perception (Blakemore et al., 1999a; Kilteni and Ehrsson, 2017a). Alternatively, the functional connectivity observed could represent somatosensory input conveyed from the cortex to the cerebellum. It was recently suggested that the cerebellar BOLD signal might primarily represent the activity of granule cells, mossy fibers or parallel fibers (Diedrichsen et al., 2010), and not changes in the spike rate of Purkinje cells (i.e., the cells that are typically presumed to encode prediction errors (Ishikawa et al., 2016)) or climbing fiber activity (Schlerf et al., 2012) (which shows characteristics suitable for computing the prediction error signal (Ishikawa et al., 2016)). Mossy fiber input could originate in the neocortex and be conveyed to the cerebellum via the pontine nuclei (Diedrichsen and Bastian, 2013). According to this interpretation, somatosensory areas project to the cerebellum to convey the received tactile feedback that could be used for computing the prediction error, for example, by contrasting the received with the predicted feedback. A third interpretation, motivated by seminal animal tracing studies, would be the case of a reciprocal exchange of information between the cerebellum and the cortex. Using both retrograde and anterograde virus injections, Kelly and Strick (2003) demonstrated the existence of closed cerebrocerebellar loops (Bostan et al., 2013); Purkinje cells located primarily at lobules IV, V and VI project to the monkey arm area in M1 and conversely, M1 projects to granule cells located primarily at Lobules IV, V and VI. Accordingly, the functional connectivity observed in our study could indicate a closed cerebrocerebellar loop between the cerebellum and the sensory cortex, in which the cerebellum sends a cancelation signal to somatosensory areas and the somatosensory areas send back tactile feedback to properly update the internal forward models. Finally, although functional connectivity does not necessarily reflect structural connectivity (Eickhoff and Müller, 2015), in our study, cerebellar regions showed correlated activity with sensorimotor areas that are predicted by earlier monkey anatomical tracing studies (Kelly and Strick, 2003; Lu et al., 2007), which might suggest that the functional connectivity effect we observed is related to anatomical connections between the involved regions. Indeed, the observed task-related functional connectivity pattern is consistent with recent findings in resting state data describing spontaneous functional couplings between lobules VI/Crus I and inferior parietal lobule and between lobule VI and the postcentral gyrus (Bernard et al., 2012), which are indicative of underlying anatomical pathways between these structures.

The cerebellar areas activated or changing connectivity strength in the present study were localized mainly in the posterior part of lobule VI, at its border with lobule Crus I and at lobule Crus I (**Figures 3-5**, **Figure 3-3**, **Figure 4-1**, **Figure 5-7**, **Figure 5-9**, **Figure 5-12**). Lobule VI is part of the primary sensorimotor body representation in the cerebellum, while lobules VII/VIII constitute the second sensorimotor representation (Grodd et al., 2001; Diedrichsen et al., 2005; Stoodley and Schmahmann, 2009; O’Reilly et al., 2010; Buckner et al., 2011; Bostan et al., 2013; Guell et al., 2018; King et al., 2018). Influential animal studies have provided evidence for a direct anatomical connection between lobules IV, V, VI and Crus I and motor cortical regions (Kelly and Strick, 2003; Lu et al., 2007; Bostan et al., 2013). Similarly, in humans, resting state data analysis showed strong functional connectivity between lobule VI and the contralateral motor cortex (Krienen and Buckner, 2009; O’Reilly et al., 2010; Bernard et al., 2012). Moreover, lobule VI has been shown to be part of the so-called “intrinsic connectivity sensorimotor network” (Habas et al., 2009) to exhibit the strongest correlation with the somatosensory and the motor cortex among other cerebellar areas (O’Reilly et al., 2010), to have strong functional connections with cerebral networks related to premotor cortex and supplementary motor area (Buckner et al., 2011) and to represent sensorimotor prediction errors (Schlerf et al., 2012).

However, while the anterior part of cerebellar lobule VI – the part adjacent to the primary fissure – is considered to be involved in sensorimotor functions, both the posterior part of lobule VI and lobule Crus I are thought to be involved in cognitive processes (Diedrichsen and Bastian, 2013; Baumann et al., 2015; Sokolov et al., 2017; Guell et al., 2018; Schmahmann, 2018). Why does somatosensory attenuation recruit cerebellar areas that are not traditionally considered related to sensorimotor function? Given the purely sensorimotor nature of our task (i.e., pressing the finger against the sensor and feeling the touch) and the fact that the corticocerebellar connectivity was modulated by the participants’ behavioral attenuation, it is highly unlikely that these cerebellar effects are driven by the participants engaging in cognitive processes during the experiment, including the rest condition (King et al., 2018). In contrast, these cerebellar effects speak in favor of a process finely tuned to the attenuation phenomenon. Schlerf et al. (2010) proposed the existence of a third sensorimotor representation in lobule VI after observing prominent activation in lobules VI/Crus I when participants performed complex (but not simple) movements with their fingers or toes. However, we consider this interpretation highly unlikely, since the pressing movements required in the tasks of the present study cannot be considered either complex or requiring any special motor coordination.

Alternatively, the cerebellar areas in the posterior cerebellum could, in addition to or in collaboration with the areas corresponding to the first and second sensorimotor representations, be involved in the predictive attenuation of self-generated input. According to this view, those posterior areas could act as intracerebellar units that process input conveyed from the sensorimotor anterior (V-VI) and/or posterior (VII/VIII) arm representations. It is quite noteworthy that findings in the literature support this view: Blakemore et al. (1998) found that the peak cerebellar activation observed when contrasting self to externally generated touch was localized for three subjects in lobule VI and for the other three in lobule Crus I (Blakemore et al., 1999b). These posterior cerebellar peaks were functionally coupled with somatosensory areas (Blakemore et al., 1999b). Moreover, when delays were introduced between movement of the right hand and somatosensory feedback on the left hand, the cerebellar areas that regressed on these sensory prediction errors elicited by the varying degrees of asynchrony were indeed observed to be situated in lobules VI and Crus I (Blakemore et al., 2001). In a study by Imamizu and colleagues (Imamizu et al., 2000), the learning of a new tool – viewed as the learning of a new internal model – was reflected in activity “*near the posterior superior fissure*”, i.e., the fissure that separates lobule VI from lobule Crus I. Additionally, cerebellar patients with lesions in Crus I have been shown to present disturbed adaptation to reaches with visuomotor and forcefield perturbations (Donchin et al., 2012) – tasks that require learning through prediction errors – and PET imaging has revealed Crus I/Crus II activation in a visuomotor perturbation task with healthy participants as well (Krakauer, 2003). Furthermore, our findings that the posterior cerebellum (VI/Crus I) increased its connectivity with the anterior part of VI (primary sensorimotor representation) and the posterior lobules VII/VIII (secondary sensorimotor representation) further support this view. Indeed, the medial location of the peaks within VII/VIII in our study is consistent with the medial representation of the hands within lobule VIII (Grodd et al., 2001; King et al., 2018), while the lateral location of the peaks within VI is in agreement with the lateral representation of the hands within lobules V/VI (Grodd et al., 2001). Both couplings (posterior VI and VIII, posterior VI/Crus I and anterior VI) are consistent with previous resting-state data that showed significant spontaneous functional correlations between Crus I and the anterior cerebellum, as well as between lobule VI with VIIb and VIIIa (Bernard et al., 2012). When further considering that our functional connectivity patterns were stronger the more participants attenuated their self-generated forces, we speculate that posterior VI/Crus acts as an intracerebellar hub that computes the prediction of self-generated information using sensory and motor information about the two hands that is conveyed from the traditional sensorimotor representations in the cerebellum that are interconnected with the sensorimotor cortex.

Sensory attenuation has been proposed to be an effective mechanism serving self-other distinction (Blakemore et al., 2000; Blakemore and Frith, 2003). Our findings suggest that corticocerebellar functional connectivity implements the sensory attenuation phenomenon and that the strength of this connection predicts the degree of sensory attenuation observed behaviorally across individuals. It is then logical to anticipate that people exhibiting reduced somatosensory attenuation would have reduced functional corticocerebellar connectivity and experience a more imprecise distinction between the self and the external world. In this context, it is interesting to note that schizophrenic patients are observed to misattribute self-generated input to external causes (auditory hallucinations, delusions of control) (Fletcher and Frith, 2009); additionally, they show reduced corticocerebellar functional connectivity (Collin et al., 2011; Repovs et al., 2011) and attenuate their self-generated touches to a weaker degree compared to healthy controls as measured in the force-matching task (Shergill et al., 2005, 2014). Finally, somatosensory attenuation has been used as an explanation for why people cannot tickle themselves (Weiskrantz et al., 1971; Blakemore et al., 2000). Speculatively, our results could thus be informative about the neural mechanism of ticklishness, and we hypothesize that disruption of corticocerebellar functional connectivity in healthy subjects by means of transcranial magnetic stimulation could make self-generated touch feel more intense and ticklish.

## Acknowledgments

We thank Christian Houborg for collecting the force-matching behavioral data, Paul Rousse for his technical support during the scans and Rouslan Sitnikov for assisting in the signal-to-noise measurements. Konstantina Kilteni was supported by the Marie Skłodowska-Curie Intra-European Individual Fellowship (#704438). The project was funded by the Swedish Research Council, Torsten Söderbergs Stiftelse, and Göran Gustafssons Stiftelse.

