## Supplementary figures and images for "Functional connectivity between the cerebellum and somatosensory areas implements the attenuation of self-generated touch"

### Figure 1-1

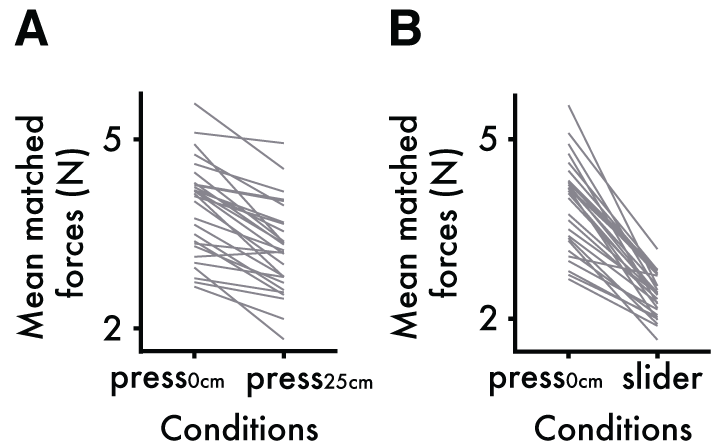

### Figure 2-1

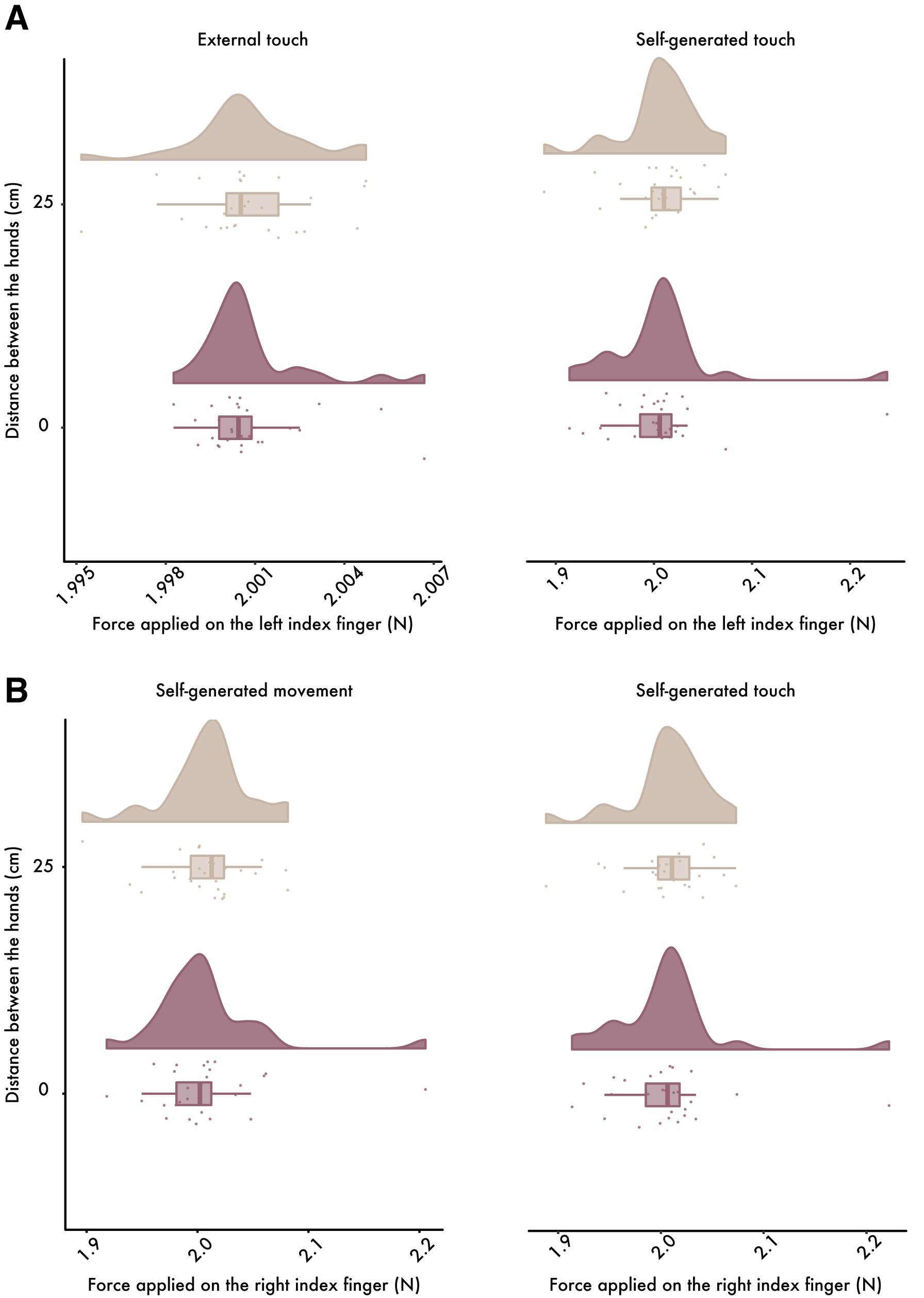

### Figure 3-1

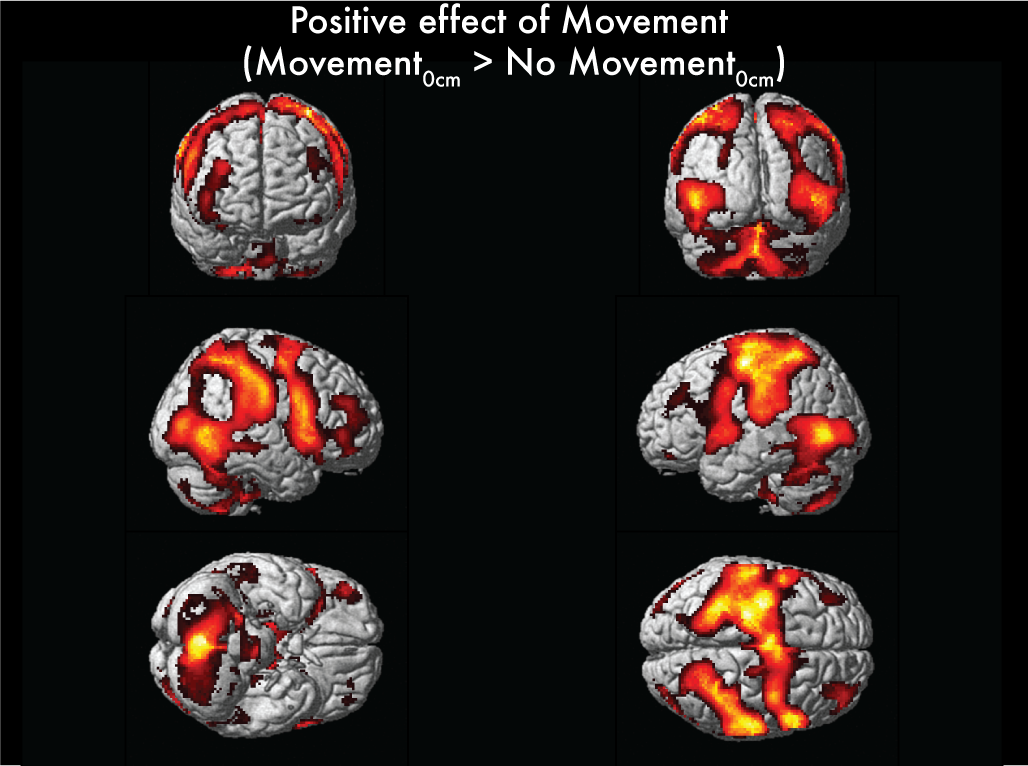

### Figure 3-2

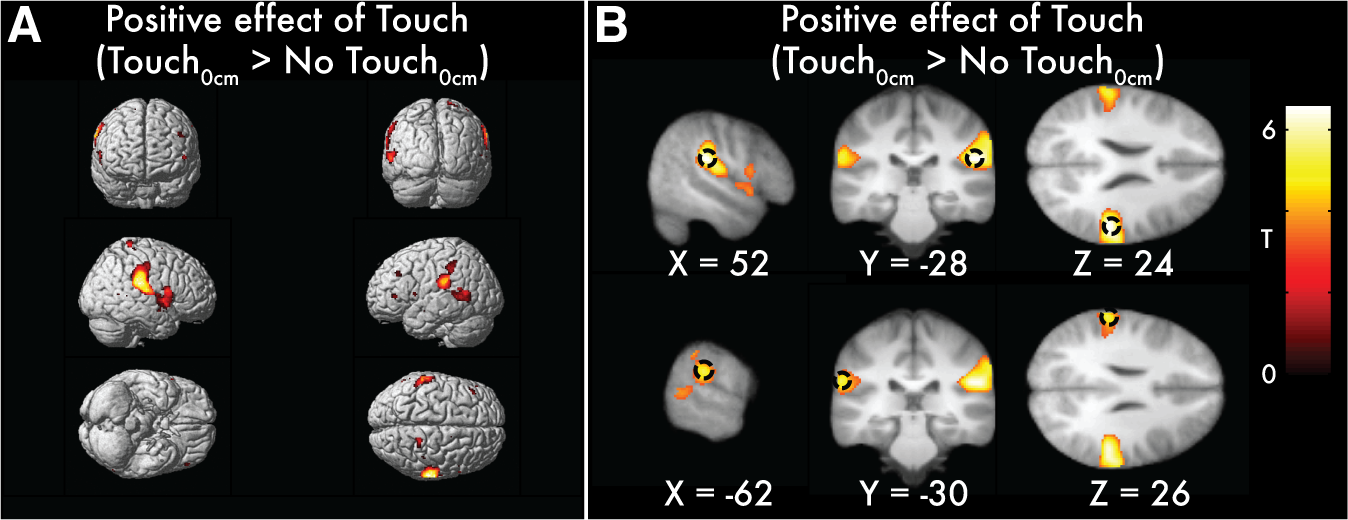

### Figure 3-3

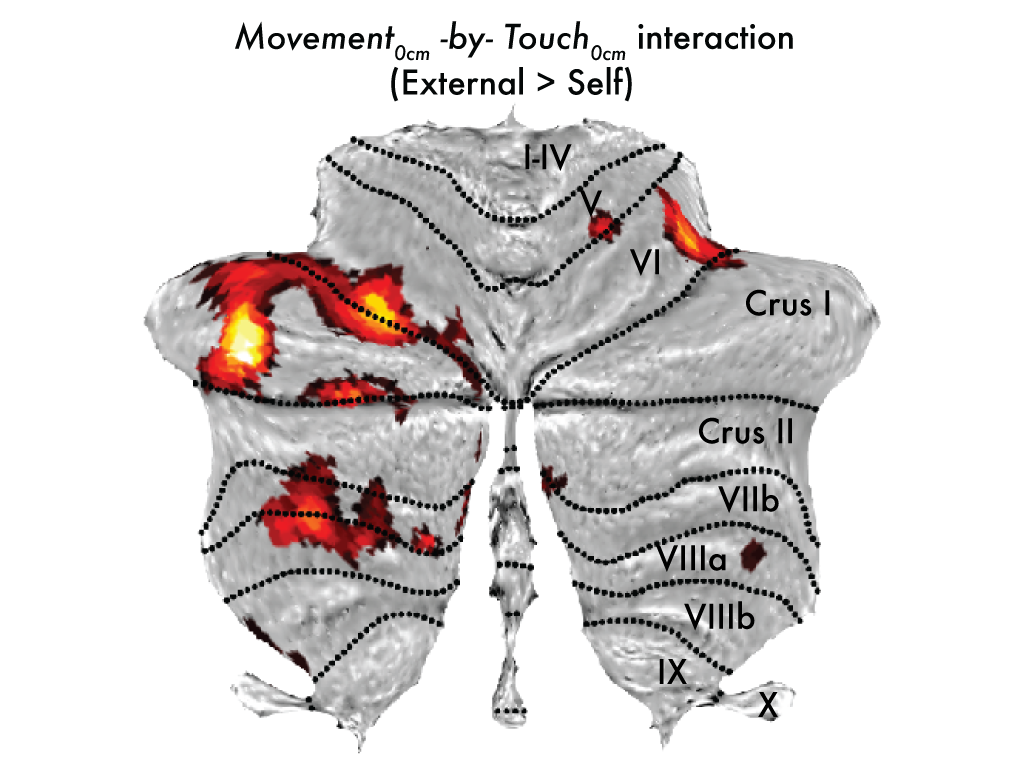

### Figure 4-1

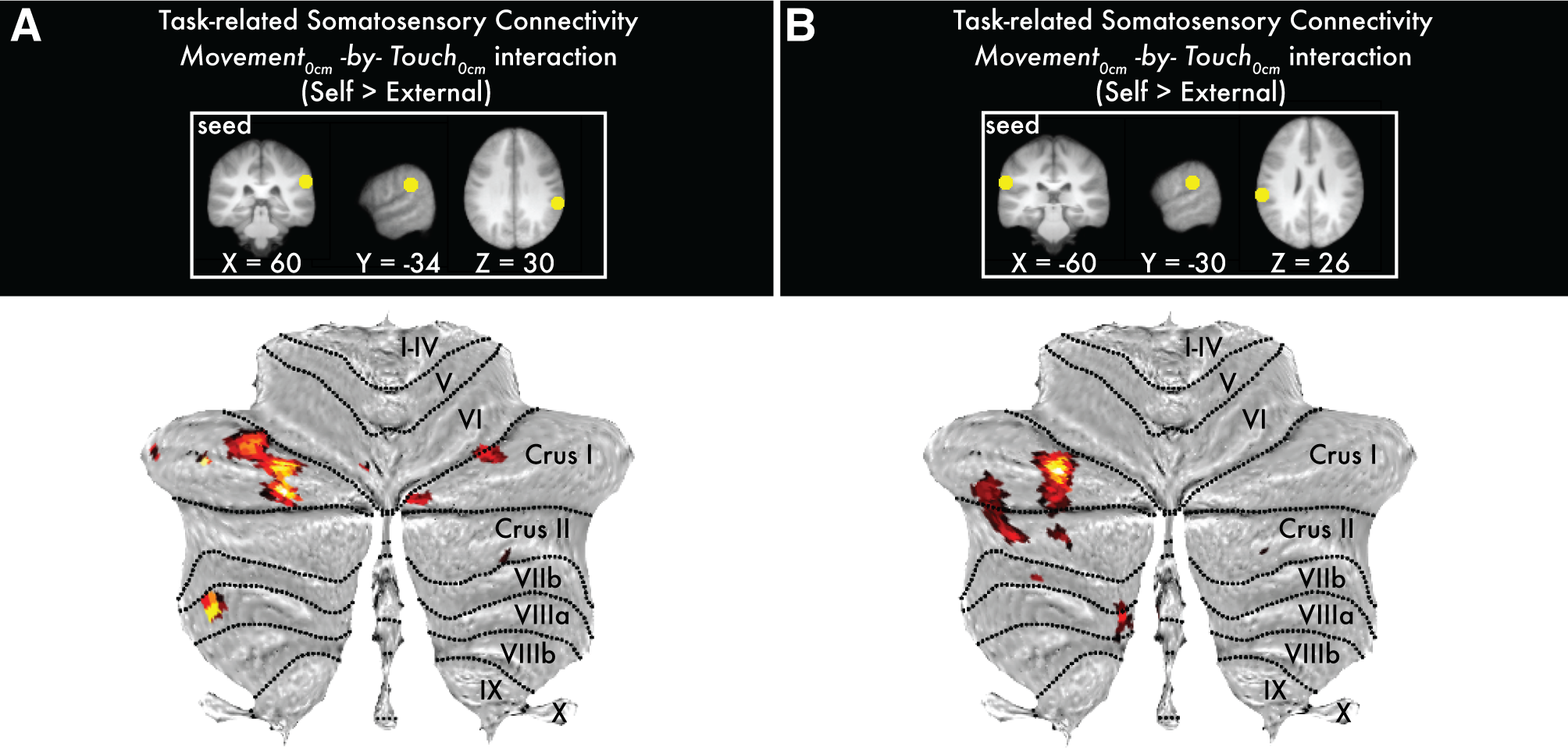

### Figure 4-2

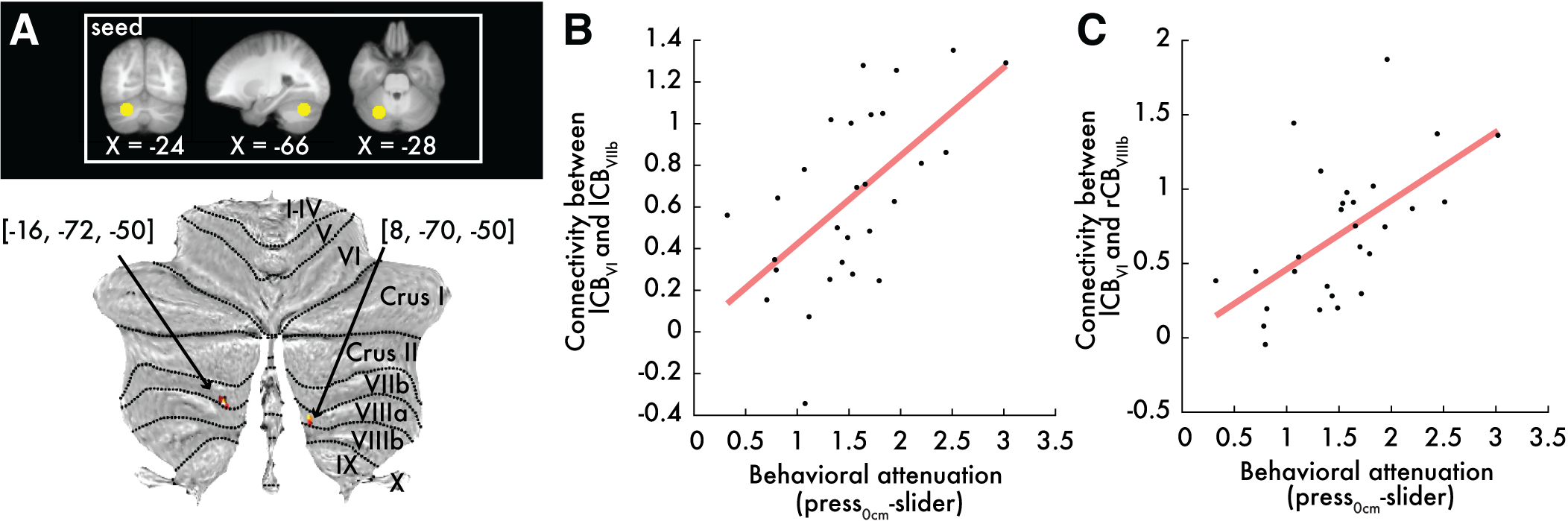

### Figure 5-1

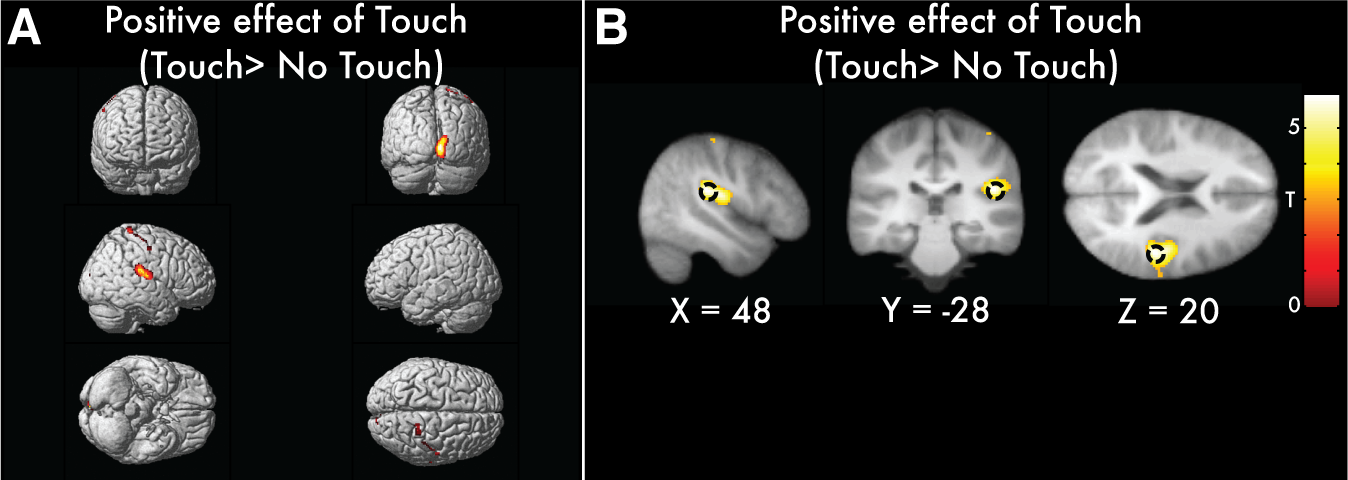

### Figure 5-3

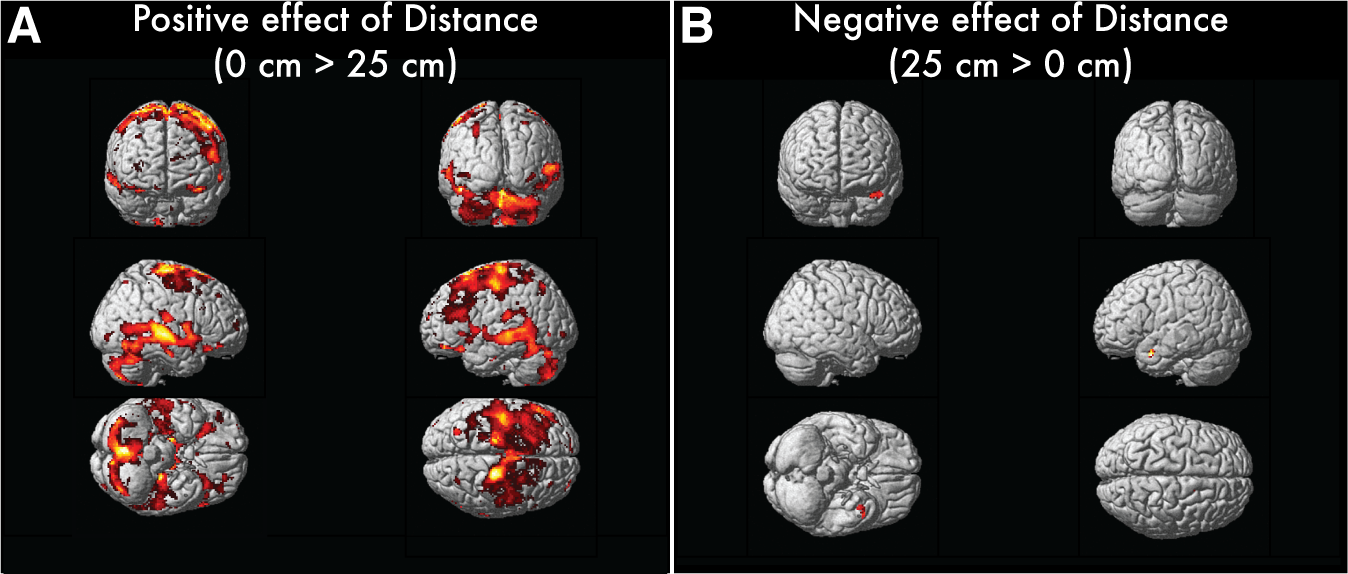

### Figure 5-7

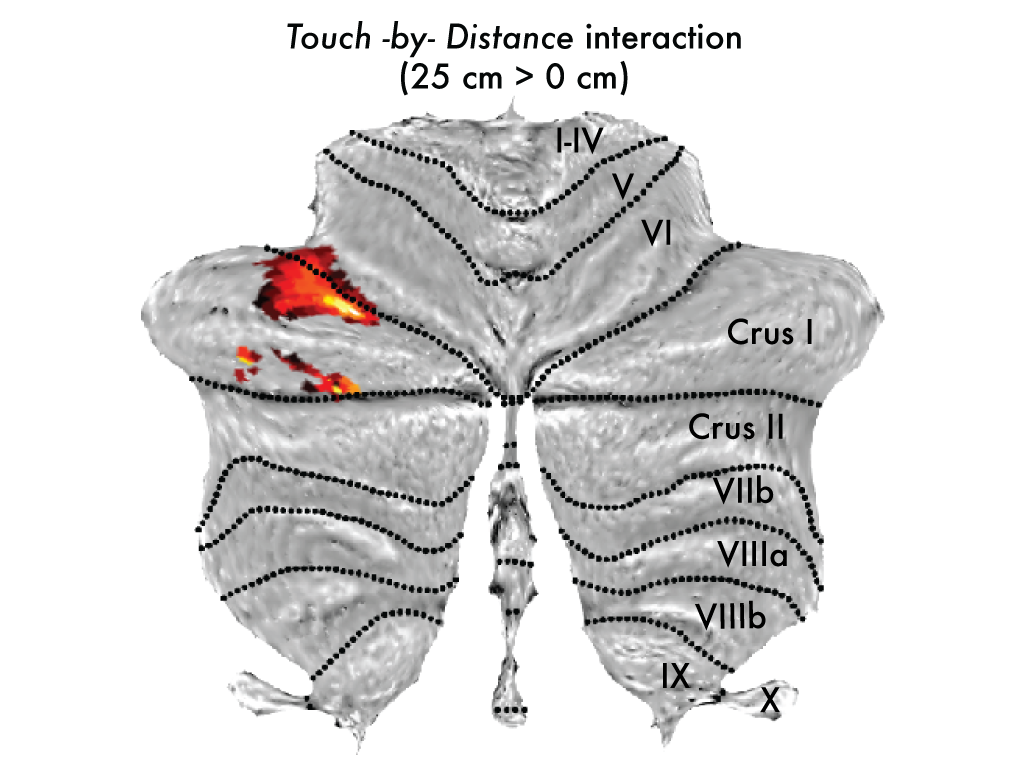

### Figure 5-9

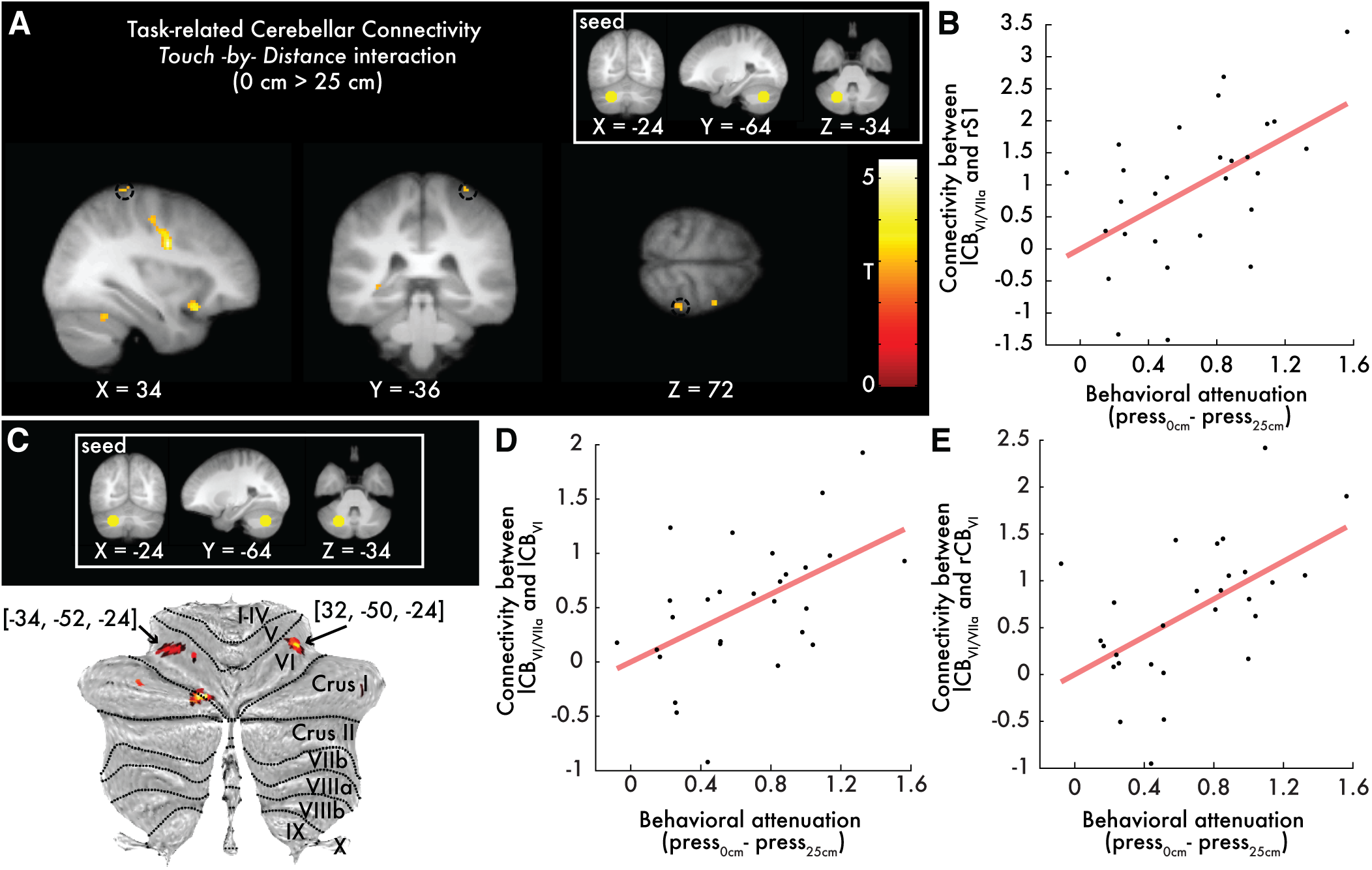

### Figure 5-12

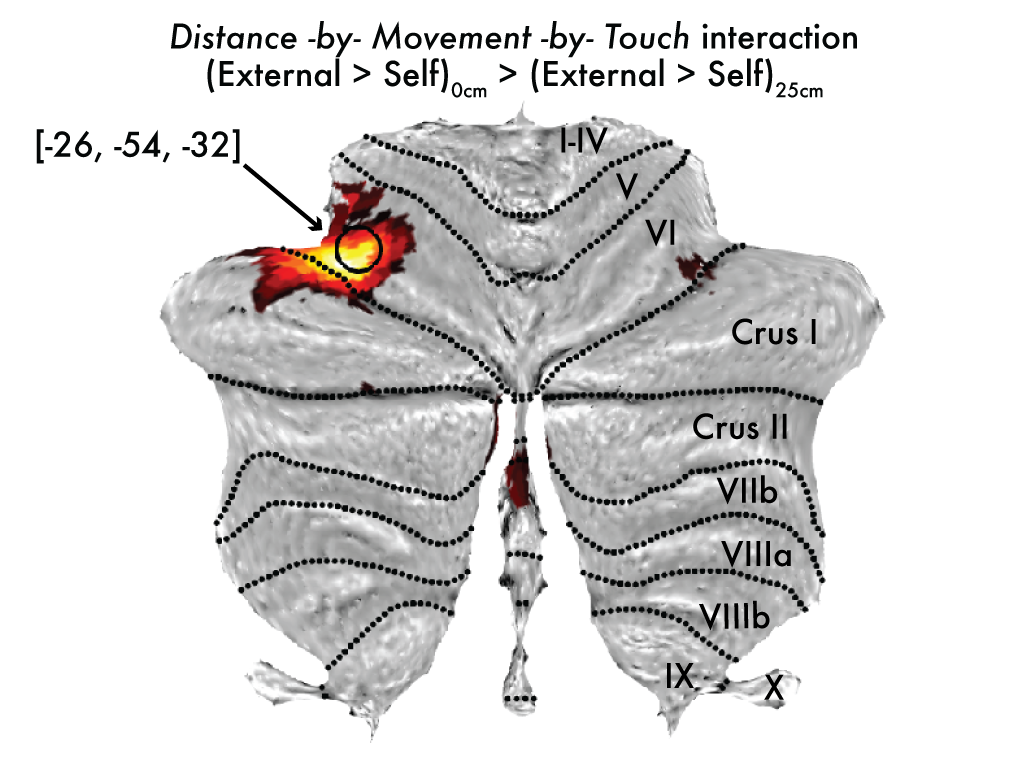
