## Supplementary material for "Functional connectivity between the cerebellum and somatosensory areas implements the attenuation of self-generated touch": Figure 5-2

**Extended Figure 5-2. Activation peaks for the positive effect of touch.** Peaks reflecting greater effects during tactile stimulation of the left index finger compared to no stimulation (Touch > No Touch).

| Brain region | Cluster size (voxels) | MNI coordinates (mm) | | | *z* | *p* |
| --- | --- | --- | --- | --- | --- | --- |
|  |  | x | y | z |  |  |
| R calcarine sulcus | 657 | 10 | -88 | 6 | 5.48 | *p* = 0.001 FWE-corrected |
| R lingual gyrus |  | 10 | -82 | -6 | 5.03 | *p* = 0.006 FWE-corrected |
| R parietal operculum | 766 | 48 | -28 | 20 | 5.44 | *p* = 0.001 FWE-corrected |
| R parietal operculum |  | 46 | -18 | 18 | 5.36 | *p* = 0.001 FWE-corrected |
| R supramarginal gyrus |  | 66 | -22 | 18 | 3.15 | *p* < 0.001 uncorrected |
| R superior parietal gyrus | 53 | 20 | -40 | 76 | 3.84 | *p* < 0.001 uncorrected |
| R hippocampus | 5 | 20 | -6 | -14 | 3.40 | *p* < 0.001 uncorrected |
| R postcentral gyrus (S1) | 14 | 56 | -14 | 50 | 3.37 | *p* < 0.001 uncorrected |
| R postcentral gyrus (S1) | 17 | 44 | -28 | 66 | 3.35 | *p* < 0.001 uncorrected |
| R postcentral gyrus (S1) |  | 50 | -20 | 60 | 3.33 | *p* < 0.001 uncorrected |
