## Supplementary material for "Functional connectivity between the cerebellum and somatosensory areas implements the attenuation of self-generated touch": Figure 5-4

**Extended Figure 5-4 Activation peaks for the positive effect of distance.** Peaks reflecting greater effects when the hands had no horizontal distance compared to when the hands were spatially separated (0 cm > 25 cm). We report a maximum of 3 peaks per cluster and only the peaks that survived FWE correction (*p* < 0.05) for spatial restrictions.

| Brain region | Cluster size (voxels) | MNI coordinates (mm) | | | *z* | *p* |
| --- | --- | --- | --- | --- | --- | --- |
|  |  | x | y | z |  |  |
| L fusiform gyrus | 3658 | -42 | -46 | -20 | 6.77 | *p* <0.001 FWE-corrected |
| L hippocampus |  | -26 | -30 | -10 | 6.19 | *p* <0.001 FWE-corrected |
| R globus pallidus |  | 26 | -10 | -10 | 5.98 | *p* <0.001 FWE-corrected |
| R cerebellum IX | 77 | 8 | -58 | -58 | 6.43 | *p* <0.001 FWE-corrected |
| L cerebellum IX |  | -6 | -58 | -58 | 5.55 | *p* <0.001 FWE-corrected |
| R precentral gyrus (M1/PMd) | 226 | 22 | -18 | 76 | 6.11 | *p* <0.001 FWE-corrected |
| R precentral gyrus (M1) |  | 20 | -24 | 66 | 5.39 | *p* =0.001 FWE-corrected |
| R superior frontal gyrus |  | 10 | -6 | 76 | 4.71 | *p* =0.023 FWE-corrected |
| L inferior frontal gyrus | 52 | -54 | 26 | 24 | 6.05 | *p* <0.001 FWE-corrected |
| L inferior frontal gyrus |  | -50 | 34 | 22 | 5.07 | *p* =0.005 FWE-corrected |
| L precentral gyrus (M1/PMd) | 276 | -44 | -10 | 60 | 6.01 | *p* <0.001 FWE-corrected |
| L precentral gyrus (PMd) |  | -18 | -18 | 78 | 5.98 | *p* <0.001 FWE-corrected |
| L postcentral gyrus (S1) |  | -46 | -20 | 62 | 5.62 | *p* <0.001 FWE-corrected |
| L superior frontal gyrus | 199 | -18 | 20 | 66 | 6.00 | *p* <0.001 FWE-corrected |
| L middle frontal gyrus |  | -26 | 30 | 54 | 5.87 | *p* <0.001 FWE-corrected |
| L middle frontal gyrus |  | -38 | 16 | 56 | 5.66 | *p* <0.001 FWE-corrected |
| R fusiform gyrus | 143 | 42 | -48 | -18 | 5.59 | *p* <0.001 FWE-corrected |
| R superior temporal sulcus | 275 | 62 | -20 | -2 | 5.46 | *p* =0.001 FWE-corrected |
| R superior temporal sulcus |  | 48 | -16 | -10 | 5.38 | *p* =0.001 FWE-corrected |
| R superior temporal gyrus |  | 54 | -26 | 2 | 5.11 | *p* =0.004 FWE-corrected |
| R cerebellum VIIa Crus II | 25 | 36 | -72 | -52 | 5.31 | *p* =0.001 FWE-corrected |
| R cerebellum VIIa Crus I | 25 | 44 | -62 | -34 | 5.24 | *p* =0.002 FWE-corrected |
| L cerebellum VIIb | 28 | -32 | -74 | -52 | 5.06 | *p* =0.005 FWE-corrected |
| L cerebellum VIIa Crus II |  | -18 | -80 | -48 | 4.91 | *p* =0.010 FWE-corrected |
| L middle temporal sulcus | 90 | -52 | -20 | -8 | 5.05 | *p* =0.005 FWE-corrected |
| L middle temporal sulcus |  | -54 | -28 | -6 | 4.90 | *p* =0.010 FWE-corrected |
| R cerebellum VIIa Crus I | 18 | 34 | -80 | -38 | 5.05 | *p* =0.005 FWE-corrected |
| R inferior frontal gyrus | 9 | 30 | 26 | -20 | 4.85 | *p* =0.013 FWE-corrected |
| R middle temporal gyrus | 4 | 58 | -38 | -10 | 4.76 | *p* =0.019 FWE-corrected |
| R superior temporal gyrus | 9 | 54 | 8 | -8 | 4.67 | *p* =0.027 FWE-corrected |
| L middle frontal gyrus | 4 | -42 | 32 | 38 | 4.63 | *p* =0.032 FWE-corrected |
| R superior temporal gyrus | 5 | 66 | -34 | 6 | 4.63 | *p* =0.032 FWE-corrected |
| L parietal operculum | 5 | -44 | -30 | 20 | 4.61 | *p* =0.035 FWE-corrected |
| R cingulate sulcus (CMA) | 4 | 8 | -16 | 48 | 4.56 | *p* =0.043 FWE-corrected |
