## Supplementary material for "Functional connectivity between the cerebellum and somatosensory areas implements the attenuation of self-generated touch": Figure 5-5

**Extended Figure 5-5. Activation peaks for the negative effect of distance.** Peaks reflecting greater effects when the hands were spatially separated compared to when the hands had no horizontal distance (25 cm > 0 cm).

| Brain region | Cluster size (voxels) | MNI coordinates (mm) | | | Z value | P value |
| --- | --- | --- | --- | --- | --- | --- |
|  |  | x | y | z |  |  |
| R thalamus | 26 | 12 | -16 | 18 | 4.05 | *p* < 0.001 uncorrected |
| L thalamus | 127 | -16 | -26 | 16 | 3.58 | *p* < 0.001 uncorrected |
| L collateral sulcus | 12 | -30 | -48 | -2 | 3.51 | *p* < 0.001 uncorrected |
| L middle temporal gyrus | 81 | -46 | 6 | -24 | 3.35 | *p* < 0.001 uncorrected |
