## Supplementary material for "Functional connectivity between the cerebellum and somatosensory areas implements the attenuation of self-generated touch": Figure 5-6

**Extended Figure 5-6. Activation peaks for the *Touch -by- Distance* interaction.** Peaks reflecting greater effects during the simultaneous presentation of movement and touch compared to the self-generated touch condition (Direction: 25 cm > 0 cm) at an uncorrected threshold of *p* < 0.001.

| Brain region | Cluster size (voxels) | MNI coordinates (mm) | | | *z* | *p* |
| --- | --- | --- | --- | --- | --- | --- |
|  |  | x | y | z |  |  |
| L cerebellum VIIa Crus I/ VI | 12 | -24 | -64 | -34 | 3.42 | *p* = 0.034 FWE-corrected^*1^ |
| R inferior part of precentral sulcus | 4 | 34 | 2 | 28 | 3.23 | *p* < 0.001 uncorrected |

^1^ Uncorrected cluster size is 16 and changed to 12 after small volume correction.
