## Supplementary material for "Functional connectivity between the cerebellum and somatosensory areas implements the attenuation of self-generated touch": Figure 5-8

**Extended Figure 5-8. Cerebellar activation peaks for the *Touch -by- Distance* interaction at a lower statistical threshold.** Peaks reflecting greater effects when the touch is presented in the context of a 25 cm hands’ distance compared to when it is presented in the absence of distance at an uncorrected threshold of *p* < 0.005.

| Brain region | Cluster size (voxels) | MNI coordinates (mm) | | | *z* | *p* |
| --- | --- | --- | --- | --- | --- | --- |
|  |  | x | y | z |  |  |
| L cerebellum VIIa Crus I /VI | 212 | -24 | -64 | -34 | 3.42 | *p* < 0.001 uncorrected^1^ |
| L cerebellum VIIa Crus I /VI | 143 | -24 | -64 | -34 | 3.42 | *p* < 0.001 uncorrected^1^ |
| L cerebellum VIIa Crus I |  | -34 | -60 | -32 | 3.00 | *p* = 0.001 uncorrected |
| L cerebellum VIIa Crus I | 5 | -46 | -62 | -38 | 3.16 | *p* = 0.001 uncorrected |

^1^ Different cluster sizes. When applying the anatomical mask for the entire cerebellum, the reported cluster size is 143.
