## Supplementary material for "Functional connectivity between the cerebellum and somatosensory areas implements the attenuation of self-generated touch": Figure 5-10

**Extended Figure 5-10. Peaks that increased their connectivity with the left cerebellum as a function of behavioral attenuation** **at a lower statistical threshold.** Peaks reflecting greater connectivity with the left cerebellar seed when the touch is presented in the absence of hands’ distance compared to when it is presented in the context of a 25 cm distance in relation to force-matching task performance (*Touch -by- Distance* interaction, Direction: 0 cm > 25 cm) at an uncorrected threshold of *p* < 0.005.

| Brain region | Cluster size (voxels) | MNI coordinates (mm) | | | *z* | *p* |
| --- | --- | --- | --- | --- | --- | --- |
|  |  | x | y | z |  |  |
| R inferior frontal gyrus, pars orbitalis | 114 | 38 | 20 | -18 | 4.40 | *p* < 0.001 uncorrected |
| R precentral gyrus | 213 | 34 | -14 | 50 | 2.90 | *p* = 0.002 uncorrected |
| L middle occipital gyrus | 202 | -42 | -64 | -2 | 3.82 | *p* < 0.001 uncorrected |
| L fusiform gyrus |  | -42 | -60 | -12 | 2.87 | *p* = 0.002 uncorrected |
| L fusiform gyrus |  | -32 | -54 | -18 | 2.87 | *p* = 0.002 uncorrected |
| L cerebellum VI |  | -34 | -52 | -24 | 2.83 | *p* = 0.002 uncorrected |
| L precuneus | 58 | -10 | -66 | 40 | 3.43 | *p* < 0.001 uncorrected |
| L middle frontal gyrus | 26 | -26 | 32 | 32 | 3.36 | *p* < 0.001 uncorrected |
| R subcentral gyrus | 32 | 48 | 2 | 14 | 3.28 | *p* = 0.001 uncorrected |
| L forth occipital gyrus | 113 | -20 | -80 | -18 | 3.21 | *p* = 0.001 uncorrected |
| L forth occipital gyrus |  | -28 | -80 | -8 | 3.05 | *p* = 0.001 uncorrected |
| R cingulate gyrus | 59 | 8 | -4 | 40 | 2.88 | *p* = 0.002 uncorrected |
| R superior temporal gyrus | 20 | 48 | -4 | -12 | 3.07 | *p* = 0.001 uncorrected |
| R precentral gyrus | 6 | 32 | -10 | 74 | 3.07 | *p* = 0.001 uncorrected |
| R cerebellum VI | 21 | 32 | -50 | -24 | 3.05 | *p* = 0.001 uncorrected |
| R superior parietal gyrus | 9 | 8 | -68 | 66 | 2.97 | *p* = 0.001 uncorrected |
| R superior temporal gyrus | 30 | 58 | 0 | 0 | 2.91 | *p* = 0.002 uncorrected |
| L fusiform gyrus/cerebellum | 7 | -46 | -66 | -22 | 2.89 | *p* = 0.002 uncorrected |
| R postcentral gyrus | 11 | 34 | -36 | 72 | 2.86 | *p* = 0.002 uncorrected |
| L postcentral gyrus (posterior part of the crown) | 12 | -68 | -18 | 32 | 2.82 | *p* = 0.002 uncorrected |
| L calcarine sulcus | 15 | -24 | -64 | 4 | 2.78 | *p* = 0.003 uncorrected |
| L central operculum | 4 | -50 | 2 | 2 | 2.78 | *p* = 0.003 uncorrected |
| R superior temporal gyrus | 7 | 68 | -32 | 12 | 2.72 | *p* = 0.003 uncorrected |
| L hippocampus | 5 | -34 | -36 | -2 | 2.71 | *p* = 0.003 uncorrected |
| R superior temporal gyrus | 9 | 44 | -14 | -6 | 2.66 | *p* = 0.004 uncorrected |
