## Supplementary material for "Functional connectivity between the cerebellum and somatosensory areas implements the attenuation of self-generated touch": Figure 5-11

**Extended Figure 5-11. Peaks that increased their connectivity with the left cerebellum at a lower statistical threshold without the participants’ attenuation covariate.** Peaks reflecting greater connectivity with the left cerebellar seed when the touch is presented in the absence of hands’ distance compared to when it is presented in the context of a 25 cm distance (*Touch -by- Distance* interaction, Direction: 0 cm > 25 cm) at an uncorrected threshold of *p* < 0.005.

| Brain region | Cluster size (voxels) | MNI coordinates (mm) | | | *z* | *p* |
| --- | --- | --- | --- | --- | --- | --- |
|  |  | x | y | z |  |  |
| L gyrus rectus | 31 | -2 | 52 | -28 | 3.49 | *p* < 0.001 uncorrected |
| L cerebellum VIIa Crus II | 6 | -12 | -92 | -40 | 3.39 | *p* < 0.001 uncorrected |
| L cerebellum IX | 46 | -2 | -60 | -48 | 3.33 | *p* < 0.001 uncorrected |
| L superior frontal gyrus | 176 | -18 | 58 | 22 | 3.22 | *p* = 0.001 uncorrected |
| L superior frontal gyrus |  | -20 | 56 | 36 | 2.70 | *p* = 0.004 uncorrected |
| L hippocampus | 19 | -32 | -36 | -8 | 3.10 | *p* = 0.001 uncorrected |
| R gyrus rectus | 4 | 6 | 14 | -24 | 3.06 | *p* = 0.001 uncorrected |
| L superior frontal gyrus | 20 | -14 | 20 | 52 | 3.00 | *p* = 0.001 uncorrected |
| L inferior occipital gyrus | 14 | -16 | -102 | -16 | 2.96 | *p* = 0.002 uncorrected |
| L parahippocampal gyrus | 15 | -28 | -26 | -24 | 2.92 | *p* = 0.002 uncorrected |
| L frontomarginal gyrus | 8 | -14 | 60 | -18 | 2.90 | *p* = 0.002 uncorrected |
| L caudate nucleus | 26 | -8 | 16 | 6 | 2.87 | *p* = 0.002 uncorrected |
| L superior temporal sulcus | 7 | -48 | -12 | -18 | 2.76 | *p* = 0.003 uncorrected |
| L precuneus | 10 | -16 | -58 | 26 | 2.73 | *p* = 0.003 uncorrected |
