## Supplementary material for "Functional connectivity between the cerebellum and somatosensory areas implements the attenuation of self-generated touch": Table 2-1

**Extended Table 2-1. Activation peaks for the positive effect of movement.** Peaks reflecting greater effects during movement of the right index finger compared to rest (Movement_0cm_ > No Movement_0cm_). Only the peaks that survived the FWE correction (*p* < 0.05) are reported for spatial restrictions.

| Brain region | Cluster size (voxels) | MNI coordinates (mm) | | | *z* | *p* |
| --- | --- | --- | --- | --- | --- | --- |
|  |  | x | y | z |  |  |
| L middle occipital gyrus | 16353 | -44 | -70 | 0 | *Inf* | *p* < 0.001 FWE-corrected |
| R middle occipital gyrus |  | 46 | -64 | 0 | *Inf* | *p* < 0.001 FWE-corrected |
| R cerebellum V |  | 10 | -50 | -16 | *Inf* | *p* < 0.001 FWE-corrected |
| R cerebellum VI |  | 24 | -50 | -26 | *Inf* | *p* < 0.001 FWE-corrected |
| R cerebellum VI |  | 4 | -62 | -24 | *Inf* | *p* < 0.001 FWE-corrected |
| R supramarginal gyrus |  | 58 | -22 | 44 | *Inf* | *p* < 0.001 FWE-corrected |
| R superior parietal gyrus |  | 22 | -60 | 62 | *Inf* | *p* < 0.001 FWE-corrected |
| R cerebellum VIIIa |  | 4 | -68 | -38 | *Inf* | *p* < 0.001 FWE-corrected |
| R intraparietal sulcus |  | 28 | -54 | 56 | *Inf* | *p* < 0.001 FWE-corrected |
| R inferior part of postcentral sulcus |  | 60 | -18 | 28 | *Inf* | *p* < 0.001 FWE-corrected |
| R cerebellum VIIIb/VIIIa |  | 8 | -70 | -46 | *Inf* | *p* < 0.001 FWE-corrected |
| R middle occipital gyrus |  | 36 | -84 | 8 | *Inf* | *p* < 0.001 FWE-corrected |
| R postcentral sulcus |  | 32 | -36 | 46 | *Inf* | *p* < 0.001 FWE-corrected |
| R cerebellum IX |  | 14 | -60 | -56 | 7.72 | *p* < 0.001 FWE-corrected |
| L cerebellum VIIIa/VIIb |  | -8 | -74 | -48 | 7.66 | *p* < 0.001 FWE-corrected |
| L cerebellum VI |  | -24 | -62 | -26 | 7.48 | *p* < 0.001 FWE-corrected |
| L precentral gyrus (M1) | 20574 | -38 | -12 | 52 | *Inf* | *p* < 0.001 FWE-corrected |
| L precentral gyrus (PMv) |  | -56 | 6 | 38 | *Inf* | *p* < 0.001 FWE-corrected |
| L precentral gyrus (PMd/M1) |  | -42 | -8 | 58 | *Inf* | *p* < 0.001 FWE-corrected |
| L anterior bank of precentral gyrus (M1) |  | -28 | -24 | 58 | *Inf* | *p* < 0.001 FWE-corrected |
| L superior frontal gyrus (SMA) |  | -6 | -8 | 54 | *Inf* | *p* < 0.001 FWE-corrected |
| L superior parietal gyrus |  | -34 | -48 | 62 | *Inf* | *p* < 0.001 FWE-corrected |
| L putamen |  | -28 | -12 | 0 | *Inf* | *p* < 0.001 FWE-corrected |
| L ventral lateral thalamic nucleus |  | -14 | -20 | 4 | *Inf* | *p* < 0.001 FWE-corrected |
| L superior parietal gyrus |  | -26 | -52 | 56 | *Inf* | *p* < 0.001 FWE-corrected |
| R inferior part of precentral sulcus |  | 54 | 12 | 18 | *Inf* | *p* < 0.001 FWE-corrected |
| R inferior part of precentral sulcus |  | 52 | 6 | 30 | *Inf* | *p* < 0.001 FWE-corrected |
| L insula |  | -42 | -2 | 8 | *Inf* | *p* < 0.001 FWE-corrected |
| L superior parietal gyrus |  | -32 | -42 | 54 | *Inf* | *p* < 0.001 FWE-corrected |
| L postcentral sulcus |  | -58 | -24 | 50 | *Inf* | *p* < 0.001 FWE-corrected |
| L postcentral sulcus |  | -56 | -24 | 36 | *Inf* | *p* < 0.001 FWE-corrected |
| R superior frontal gyrus (SMA) |  | 6 | 0 | 60 | *Inf* | *p* < 0.001 FWE-corrected |
| R cingulate sulcus | 70 | 12 | -22 | 44 | 5.69 | *p* < 0.001 FWE-corrected |
| L intraparietal sulcus | 89 | -24 | -72 | 30 | 5.44 | *p* = 0.001 FWE-corrected |
| L middle frontal gyrus | 4 | -40 | 36 | 36 | 4.94 | *p* = 0.007 FWE-corrected |
| L middle frontal gyrus | 5 | -40 | 44 | 26 | 4.91 | *p* = 0.008 FWE-corrected |
| L cerebellum IX (Hem) | 4 | -6 | -58 | -58 | 4.91 | *p* = 0.009 FWE-corrected |
| L superior temporal gyrus | 12 | -50 | 16 | -10 | 4.83 | *p* = 0.012 FWE-corrected |
