## Supplementary material for "Functional connectivity between the cerebellum and somatosensory areas implements the attenuation of self-generated touch": Table 2-2

**Extended Table 2-2. Activation peaks for the positive effect of touch.** Peaks reflecting greater effects during tactile stimulation of the left index finger compared to no stimulation (Touch_0cm_ > No Touch_0cm_).

| Brain region | Cluster size (voxels) | MNI coordinates (mm) | | | *z* | *p* |
| --- | --- | --- | --- | --- | --- | --- |
|  |  | x | y | z |  |  |
| R parietal operculum (S2) | 1796 | 52 | -28 | 24 | 6.00 | *p* < 0.001 FWE-corrected |
| R supramarginal gyrus |  | 64 | -28 | 36 | 5.09 | *p* = 0.004 FWE-corrected |
| R postcentral sulcus |  | 58 | -18 | 48 | 3.78 | *p* < 0.001 uncorrected |
| L supramarginal gyrus | 602 | -62 | -30 | 26 | 4.99 | *p* < 0.006 FWE-corrected |
| L supramarginal gyrus |  | -60 | -38 | 44 | 4.14 | *p* < 0.001 uncorrected |
| R inferior frontal gyrus, pars opercularis | 891 | 60 | 12 | 6 | 4.31 | *p* < 0.001 uncorrected |
| R insula |  | 42 | 0 | 10 | 4.25 | *p* < 0.001 uncorrected |
| R insula |  | 42 | 2 | -6 | 4.23 | *p* < 0.001 uncorrected |
| R insula |  | 42 | -10 | -12 | 3.96 | *p* < 0.001 uncorrected |
| R insula |  | 46 | -2 | 0 | 3.79 | *p* < 0.001 uncorrected |
| R inferior part of precentral sulcus |  | 50 | 10 | 14 | 3.58 | *p* < 0.001 uncorrected |
| R superior parietal gyrus | 72 | 20 | -40 | 76 | 4.26 | *p* < 0.001 uncorrected |
| L middle temporal gyrus | 253 | -60 | -50 | 6 | 3.71 | *p* < 0.001 uncorrected |
| L middle temporal gyrus |  | -64 | -44 | 10 | 3.56 | *p* < 0.001 uncorrected |
| L superior temporal sulcus |  | -52 | -50 | 14 | 3.39 | *p* < 0.001 uncorrected |
| L middle frontal gyrus | 22 | -44 | 30 | 38 | 3.57 | *p* < 0.001 uncorrected |
| L inferior frontal gyrus, pars triangularis | 23 | -50 | 38 | 6 | 3.41 | *p* < 0.001 uncorrected |
| L inferior frontal gyrus, pars opercularis | 11 | -52 | 8 | 8 | 3.40 | *p* < 0.001 uncorrected |
| R postcentral gyrus (S1) | 4 | 40 | -34 | 68 | 3.28 | *p* < 0.001 uncorrected |
| L insula | 9 | -40 | -6 | -16 | 3.20 | *p* < 0.001 uncorrected |
