## Supplementary material for "Functional connectivity between the cerebellum and somatosensory areas implements the attenuation of self-generated touch": Table 2-3

**Extended Table 2-3. Activation peaks for the *Movement_0cm_ -by- Touch_0cm_* interaction.** Peaks reflecting greater effects of touch when this is presented in the absence of movement (external) compared to when it is presented in the context of movement (self-generated) (Direction: External > Self).

| Brain region | Cluster size (voxels) | MNI coordinates (mm) | | | *z* | *p* |
| --- | --- | --- | --- | --- | --- | --- |
|  |  | x | y | z |  |  |
| R supramarginal gyrus | 1637 | 60 | -34 | 30 | 4.97 | *p* = 0.007 FWE-corrected |
| R temporal parietal junction (between the superior temporal gyrus and supramarginal gyrus) |  | 62 | -36 | 22 | 4.64 | *p* = 0.027 FWE-corrected |
| R middle temporal gyrus |  | 54 | -52 | 2 | 4.43 | *p* < 0.001 uncorrected |
| R middle temporal gyrus |  | 52 | -56 | 4 | 4.38 | *p* < 0.001 uncorrected |
| R superior temporal sulcus |  | 54 | -42 | 10 | 3.76 | *p* < 0.001 uncorrected |
| R superior temporal sulcus |  | 46 | -66 | 16 | 3.68 | *p* < 0.001 uncorrected |
| R superior frontal gyrus (pre-SMA) | 150 | 14 | 4 | 64 | 4.00 | *p* < 0.001 uncorrected |
| R superior frontal gyrus (SMA) |  | 14 | -12 | 62 | 3.45 | *p* < 0.001 uncorrected |
| L parietal operculum (S2) /supramarginal gyrus | 560 | -60 | -30 | 26 | 3.92 | *p* = 0.002 FWE-corrected^*1^ |
| L supramarginal gyrus (at the posterior bank of the postcentral sulcus) |  | -58 | -22 | 26 | 3.88 | *p* = 0.003 FWE-corrected^*^ |
| L supramarginal gyrus |  | -54 | -36 | 34 | 3.84 | *p* < 0.001 uncorrected^2^ |
| L supramarginal gyrus |  | -56 | -34 | 32 | 3.60 | *p* = 0.007 FWE-corrected^*2^ |
| L supramarginal gyrus |  | -52 | -34 | 44 | 3.74 | *p* < 0.001 uncorrected |
| L postcentral sulcus |  | -46 | -38 | 58 | 3.16 | *p* < 0.001 uncorrected |
| L precuneus | 122 | -12 | -68 | 44 | 3.90 | *p* < 0.001 uncorrected |
| L cerebellum VIIa Crus I | 30 | -44 | -60 | -36 | 3.81 | *p* < 0.001 uncorrected |
| R superior temporal sulcus | 43 | 46 | -16 | -12 | 3.80 | *p* < 0.001 uncorrected |
| R superior parietal gyrus | 203 | 14 | -52 | 66 | 3.68 | *p* < 0.001 uncorrected |
| R precuneus |  | 14 | -56 | 58 | 3.61 | *p* < 0.001 uncorrected |
| R inferior frontal gyrus, pars opercularis | 52 | 54 | 10 | 4 | 3.63 | *p* < 0.001 uncorrected |
| L precuneus | 86 | -8 | -50 | 56 | 3.60 | *p* < 0.001 uncorrected |
| L middle frontal gyrus | 73 | -32 | 42 | 20 | 3.59 | *p* < 0.001 uncorrected |
| R precentral gyrus (PM) | 51 | 48 | 4 | 42 | 3.58 | *p* < 0.001 uncorrected |
| L orbital gyrus | 43 | -32 | 46 | -16 | 3.56 | *p* < 0.001 uncorrected |
| R fusiform gyrus | 41 | 40 | -50 | -20 | 3.50 | *p* < 0.001 uncorrected |
| L insula | 56 | -36 | 2 | 6 | 3.48 | *p* < 0.001 uncorrected |
| L insula |  | -40 | 6 | 0 | 3.28 | *p* < 0.001 uncorrected |
| L cerebellum VI | 44 | -24 | -66 | -28 | 3.47 | *p* = 0.026 FWE-corrected^*3^ |
| L cerebellum VIIa Crus I |  | -24 | -64 | -36 | 3.34 | *p* < 0.001 uncorrected^4^ |
| L cerebellum VIIa Crus I |  | -22 | -64 | -34 | 3.24 | *p* < 0.001 uncorrected^4^ |
| R inferior frontal gyrus | 45 | 44 | 42 | 2 | 3.46 | *p* < 0.001 uncorrected |
| R middle frontal gyrus |  | 46 | 48 | 8 | 3.20 | *p* < 0.001 uncorrected |
| R orbital gyrus | 5 | 32 | 48 | -12 | 3.45 | *p* < 0.001 uncorrected |
| L inferior frontal gyrus, pars opercularis | 73 | -50 | 6 | 10 | 3.45 | *p* < 0.001 uncorrected |
| L cingulate gyrus or white matter | 43 | -12 | 8 | 34 | 3.34 | *p* < 0.001 uncorrected |
| L superior frontal gyrus |  | -18 | 8 | 50 | 3.20 | *p* < 0.001 uncorrected |
| R inferior frontal gyrus, pars opercularis | 12 | 42 | 18 | 6 | 3.29 | *p* < 0.001 uncorrected |
| R precuneus | 15 | 16 | -64 | 44 | 3.24 | *p* < 0.001 uncorrected |
| L cerebellum VIIb/VIIIa | 10 | -34 | -60 | -54 | 3.24 | *p* < 0.001 uncorrected |
| L middle temporal gyrus | 8 | -58 | -58 | 8 | 3.23 | *p* < 0.001 uncorrected |
| R insula | 7 | 38 | 8 | 0 | 3.22 | *p* < 0.001 uncorrected |
| R precuneus | 6 | 8 | -52 | 46 | 3.20 | *p* < 0.001 uncorrected |

**^*^** After small volume correction.

^1^ The cluster size is 560 before corrections for multiple comparisons and is reduced to 221 after small volume correction.

^2^ Repeated spatially adjacent peaks. The first peak in order is the uncorrected peak belonging to the corresponding cluster, while the second peak in order replaces the previous peak after small volume correction.

^3^ The cluster size is 52 before corrections for multiple comparisons and is reduced to 44 after small volume correction.

^4^ Repeated spatially adjacent peaks. The first peak in order is the uncorrected peak belonging to the corresponding cluster, while the second peak in order replaces the previous peak after small volume correction.
