## Supplementary material for "Functional connectivity between the cerebellum and somatosensory areas implements the attenuation of self-generated touch": Table 2-4

**Extended Table 2-4. Cerebellar activation peaks for the *Movement_0cm_ -by- Touch_0cm_* interaction at the uncorrected statistical threshold of *p* < 0.005.** Cerebellar peaks reflecting greater effects of touch when this is presented in the absence of movement (external) compared to when it is presented in the context of movement (self-generated) (Direction: External > Self) at an uncorrected threshold of *p* < 0.005.

| Brain region | Cluster size (voxels) | MNI coordinates (mm) | | | *z* | *p* |
| --- | --- | --- | --- | --- | --- | --- |
|  |  | x | y | z |  |  |
| L cerebellum VIIa Crus I | 371 | -44 | -60 | -36 | 3.81 | *p* < 0.001 uncorrected |
| L cerebellum VI |  | -24 | -66 | -28 | 3.47 | *p* < 0.001 uncorrected |
| L cerebellum VI/VIIa Crus I |  | -22 | -64 | -34 | 3.24 | *p* = 0.001 uncorrected |
| L cerebellum VIIa Crus I |  | -26 | -62 | -36 | 3.16 | *p* = 0.001 uncorrected |
| L cerebellum VI/VIIa |  | -34 | -52 | -36 | 3.13 | *p* = 0.001 uncorrected |
| L cerebellum VI |  | -12 | -66 | -26 | 2.68 | *p* = 0.004 uncorrected |
| L cerebellum VIIb /VIIIa | 205 | -34 | -60 | -54 | 3.24 | *p* = 0.001 uncorrected |
| L cerebellum VIIb |  | -12 | -76 | -50 | 3.05 | *p* = 0.001 uncorrected |
| L cerebellum VIIIa |  | -34 | -54 | -52 | 3.01 | *p* = 0.001 uncorrected |
| L cerebellum VIIIa |  | -24 | -66 | -54 | 2.96 | *p* = 0.002 uncorrected |
| L cerebellum VIIa Crus II |  | -40 | -70 | -50 | 2.84 | *p* = 0.002 uncorrected |
| L cerebellum VIIb |  | -28 | -74 | -50 | 2.80 | *p* = 0.003 uncorrected |
| L cerebellum VIIIa/VIIa Crus II | 27 | -2 | -78 | -40 | 3.01 | *p* = 0.001 uncorrected |
| Cerebellum VIIa Crus II |  | 0 | -80 | -36 | 2.86 | *p* = 0.002 uncorrected |
| R cerebellum VI | 12 | 38 | -52 | -26 | 2.92 | *p* = 0.002 uncorrected |
| R cerebellum VI | 5 | 24 | -54 | -16 | 2.83 | *p* = 0.002 uncorrected |
| L cerebellum VIIa Crus I | 8 | -26 | -80 | -34 | 2.78 | *p* = 0.003 uncorrected |
| R cerebellum VIIb/VIIIa | 5 | 34 | -52 | -52 | 2.68 | *p* = 0.004 uncorrected |
