## Supplementary material for "Functional connectivity between the cerebellum and somatosensory areas implements the attenuation of self-generated touch": Table 2-5

**Extended Table 2-5. Cerebellar activation peaks for the *Movement_0cm_ -by- Touch_0cm_* interaction.** Peaks reflecting greater effects of touch when this is presented in the absence of movement (external) compared to when it is presented in the context of movement (self-generated) (Direction: External > Self)

| Brain region | Cluster size (voxels) | MNI coordinates (mm) | | | *z* | *p* |
| --- | --- | --- | --- | --- | --- | --- |
|  |  | x | y | z |  |  |
| L cerebellum VIIa Crus I | 30 | -44 | -60 | -36 | 3.81 | *p* < 0.001 uncorrected |
| L cerebellum VI | 42 | -24 | -66 | -28 | 3.47 | *p* < 0.001 uncorrected^1^ |
| L cerebellum VI | 44 | -24 | -66 | -28 | 3.47 | *p* = 0.026 FWE-corrected^*1^ |
| L cerebellum VI/VIIa Crus I |  | -22 | -64 | -34 | 3.24 | *p* = 0.001 uncorrected |
| L cerebellum VIIb/VIIIa | 10 | -34 | -60 | -54 | 3.24 | *p* = 0.001 uncorrected |

**^*^** After small volume correction.

^1^ Different cluster sizes. When applying the anatomical mask for the entire cerebellum, the reported cluster size is 42. When correcting for small volume correction, the cluster size is 44.
