## Supplementary material for "Functional connectivity between the cerebellum and somatosensory areas implements the attenuation of self-generated touch": Table 2-6

**Extended Table 2-6. Activation peaks for the *Movement_0cm_ -by- Touch_0cm_* interaction.** Peaks reflecting greater effects of touch when this is presented in the context of movement (self-generated) compared to when it is presented in the absence of movement (external) (Direction: Self > External). Visual activations were due to the visual instructions since the messages given in the self-generated conditions (*press*, *press&feel*) were different and longer than the messages given in the externally generated conditions (*feel*, *rest*).

| Brain region | Cluster size (voxels) | MNI coordinates (mm) | | | *z* | *p* |
| --- | --- | --- | --- | --- | --- | --- |
|  |  | x | y | z |  |  |
| L middle occipital gyrus | 286 | -16 | -98 | -2 | 4.47 | *p* < 0.001 uncorrected |
| L middle occipital gyrus |  | -30 | -92 | -6 | 3.51 | *p* < 0.001 uncorrected |
| R lingual gyrus | 300 | 10 | -86 | -4 | 4.45 | *p* < 0.001 uncorrected |
