## Supplementary material for "Functional connectivity between the cerebellum and somatosensory areas implements the attenuation of self-generated touch": Table 3-1

**Extended Table 3-1. Peaks that increased their connectivity with the right supramarginal gyrus** **as a function of behavioral attenuation.** Peaks reflecting greater connectivity with the right supramarginal gyrus when the touch is presented in the context of movement (self-generated) compared to when it is presented in the absence of movement (external) in relation to force-matching task performance (*Movement_0cm_ -by- Touch_0cm_* interaction, Direction: Self > External).

| Brain region | Cluster size (voxels) | MNI coordinates (mm) | | | *z* | *p* |
| --- | --- | --- | --- | --- | --- | --- |
|  |  | x | y | z |  |  |
| R superior frontal gyrus | 43 | 18 | 56 | 24 | 4.06 | *p* < 0.001 uncorrected |
| L cingulate sulcus | 300 | -2 | 34 | 24 | 3.89 | *p* < 0.001 uncorrected |
| R cingulate sulcus |  | 12 | 30 | 26 | 3.61 | *p* < 0.001 uncorrected |
| L cingulate sulcus |  | -2 | 40 | 14 | 3.52 | *p* < 0.001 uncorrected |
| L lingual gyrus | 97 | -2 | -88 | -18 | 3.88 | *p* < 0.001 uncorrected |
| L middle frontal gyrus | 72 | -44 | 26 | 42 | 3.77 | *p* < 0.001 uncorrected |
| L cerebellum VIIa Crus I | 96 | -26 | -70 | -32 | 3.72 | *p* < 0.001 uncorrected^1^ |
| L cerebellum VIIa Crus I | 26 | -26 | -70 | -30 | 3.58 | *p* = 0.037 FWE-corrected^*1^ |
| L cerebellum VIIa Crus I |  | -34 | -62 | -30 | 3.46 | *p* < 0.001 uncorrected |
| L middle frontal gyrus | 14 | -36 | 62 | 10 | 3.68 | *p* < 0.001 uncorrected |
| L cerebellum VIIa Crus I | 5 | -50 | -42 | -38 | 3.67 | *p* < 0.001 uncorrected |
| L cerebellum VIIa Crus I | 7 | -52 | -62 | -30 | 3.64 | *p* < 0.001 uncorrected |
| R cingulate sulcus | 25 | 16 | -38 | 50 | 3.63 | *p* < 0.001 uncorrected |
| R cingulate sulcus |  | 10 | -34 | 44 | 3.27 | *p* < 0.001 uncorrected |
| R supramarginal gyrus | 14 | 62 | -38 | 42 | 3.55 | *p* < 0.001 uncorrected |
| L superior frontal gyrus | 13 | -18 | 22 | 66 | 3.48 | *p* < 0.001 uncorrected |
| R cerebellum VI/VIIa Crus I | 13 | 10 | -84 | -22 | 3.40 | *p* < 0.001 uncorrected |
| R thalamus | 5 | 12 | -24 | 16 | 3.40 | *p* < 0.001 uncorrected |
| L superior occipital gyrus | 39 | -8 | -84 | 42 | 3.38 | *p* < 0.001 uncorrected |
| R cerebellum VIIIb/IX | 11 | 16 | -38 | -52 | 3.36 | *p* < 0.001 uncorrected |
| L superior frontal gyrus | 4 | -8 | 66 | 28 | 3.32 | *p* < 0.001 uncorrected |
| R cerebellum VI | 20 | 28 | -70 | -24 | 3.31 | *p* < 0.001 uncorrected |
| R gyrus descendens | 5 | 20 | -104 | -4 | 3.31 | *p* < 0.001 uncorrected |
| R cingulate sulcus | 6 | 12 | 22 | 40 | 3.30 | *p* < 0.001 uncorrected |
| R superior frontal gyrus | 4 | 4 | 66 | 26 | 3.28 | *p* < 0.001 uncorrected |
| R lingual gyrus | 7 | 16 | -88 | -10 | 3.25 | *p* < 0.001 uncorrected |
| L supramarginal | 4 | -52 | -48 | 56 | 3.24 | *p* < 0.001 uncorrected |
| R gyrus descendens | 4 | 20 | -98 | -10 | 3.23 | *p* < 0.001 uncorrected |

**^*^** After small volume correction.

^1^ Repeated spatially adjacent peaks. The first peak in order is the uncorrected peak belonging to a cluster with a size of 96, while the second peak in order belongs to a smaller cluster with a size of 26 and replaces the previous one after small volume correction.
