## Supplementary material for "Functional connectivity between the cerebellum and somatosensory areas implements the attenuation of self-generated touch": Table 3-2

**Extended Table 3-2. Peaks that increased their connectivity with the left parietal operculum S2/SMG as a function of behavioral attenuation.** Peaks reflecting greater connectivity with the left somatosensory seed when the touch is presented in the context of movement (self-generated) compared to when it is presented in the absence of movement (external) in relation to force-matching task performance (*Movement_0cm_ -by- Touch_0cm_* interaction, Direction: Self > External).

| Brain region | Cluster size (voxels) | MNI coordinates (mm) | | | *z* | *p* |
| --- | --- | --- | --- | --- | --- | --- |
|  |  | x | y | z |  |  |
| L cerebellum VIIa Crus I | 82 | -32 | -72 | -28 | 4.09 | *p* < 0.001 uncorrected |
| L superior frontal gyrus | 348 | -8 | 66 | 26 | 4.03 | *p* < 0.001 uncorrected |
| L superior frontal gyrus |  | -2 | 68 | 10 | 3.88 | *p* < 0.001 uncorrected |
| R superior frontal gyrus |  | 2 | 64 | 22 | 3.71 | *p* < 0.001 uncorrected |
| Superior frontal gyrus |  | 0 | 50 | 8 | 3.69 | *p* < 0.001 uncorrected |
| L superior frontal gyrus |  | -4 | 58 | 14 | 3.68 | *p* < 0.001 uncorrected |
| L superior frontal gyrus |  | -6 | 62 | 8 | 3.41 | *p* < 0.001 uncorrected |
| L cingulate sulcus |  | -2 | 44 | 14 | 3.37 | *p* < 0.001 uncorrected |
| L superior occipital gyrus | 17 | -14 | -84 | 18 | 3.62 | *p* < 0.001 uncorrected |
| L superior occipital gyrus | 19 | -8 | -90 | 42 | 3.58 | *p* < 0.001 uncorrected |
| L cerebellum VIIa Crus II | 63 | -48 | -66 | -48 | 3.57 | *p* < 0.001 uncorrected |
| L cerebellum VIIa Crus I |  | -46 | -62 | -38 | 3.38 | *p* < 0.001 uncorrected |
| L cerebellum VIIIa | 11 | -10 | -68 | -42 | 3.50 | *p* < 0.001 uncorrected |
| L inferior frontal gyrus, pars triangularis | 11 | -54 | 32 | 0 | 3.42 | *p* < 0.001 uncorrected |
| L cerebellum VIIa Crus II | 11 | -36 | -76 | -42 | 3.34 | *p* < 0.001 uncorrected |
| L middle frontal gyrus | 4 | -50 | 18 | 44 | 3.17 | *p* < 0.001 uncorrected |
| L middle frontal gyrus | 5 | -42 | 18 | 40 | 3.17 | *p* < 0.001 uncorrected |
