## Supplementary material for "Functional connectivity between the cerebellum and somatosensory areas implements the attenuation of self-generated touch": Table 3-3

**Extended Table 3-3. Peaks that increased their connectivity with the left cerebellum** **as a function of behavioral attenuation.** Peaks reflecting greater connectivity with the left cerebellar seed when the touch is presented in the context of movement (self-generated) compared to when it is presented in the absence of movement (external) in relation to force-matching task performance (*Movement_0cm_ -by- Touch_0cm_* interaction, Direction: Self > External).

| Brain region | Cluster size (voxels) | MNI coordinates (mm) | | | *z* | *p* |
| --- | --- | --- | --- | --- | --- | --- |
|  |  | x | y | z |  |  |
| L caudate nucleus | 52 | -20 | 10 | 18 | 4.49 | *p* < 0.001 uncorrected |
| L middle frontal gyrus | 301 | -38 | 46 | 14 | 3.38 | *p* < 0.001 uncorrected |
| L middle frontal gyrus |  | -42 | 40 | 22 | 3.30 | *p* < 0.001 uncorrected |
| L supramarginal gyrus | 150 | -62 | -38 | 34 | 3.97 | *p* < 0.001 uncorrected^1^ |
| L supramarginal gyrus | 26 | -62 | -36 | 34 | 3.88 | *p* = 0.006 FWE-corrected^*1^ |
| L supramarginal gyrus |  | -58 | -38 | 48 | 3.50 | *p* < 0.001 uncorrected |
| R Insula | 69 | 42 | 16 | -8 | 3.90 | *p* < 0.001 uncorrected |
| L middle occipital gyrus | 6 | -30 | -92 | 26 | 3.82 | *p* < 0.001 uncorrected |
| R superior temporal gyrus | 50 | 64 | -34 | 12 | 3.78 | *p* < 0.001 uncorrected |
| L gyrus rectus | 8 | -8 | 34 | -32 | 3.63 | *p* < 0.001 uncorrected |
| R superior temporal gyrus | 12 | 26 | 16 | -42 | 3.52 | *p* < 0.001 uncorrected |
| L middle frontal gyrus | 8 | -30 | 52 | 30 | 3.49 | *p* < 0.001 uncorrected |
| L inferior frontal gyrus | 10 | -42 | 18 | -12 | 3.48 | *p* < 0.001 uncorrected |
| L middle frontal gyrus | 28 | -36 | 58 | 8 | 3.38 | *p* < 0.001 uncorrected |
| R postcentral gyrus (S1) | 14 | 36 | -32 | 70 | 3.37 | *p* = 0.028 FWE-corrected^*^ |
| L cerebellum VIIb/VIIIa | 8 | -16 | -72 | -50 | 3.36 | *p* < 0.001 uncorrected |
| R superior temporal gyrus | 7 | 64 | 4 | 0 | 3.32 | *p* < 0.001 uncorrected |
| R middle frontal gyrus | 18 | 36 | 54 | 26 | 3.32 | *p* < 0.001 uncorrected |
| R cerebellum VIIIb/VIIIa | 5 | 8 | -70 | -50 | 3.31 | *p* < 0.001 uncorrected |
| R parietal operculum (S2)/ supramarginal gyrus | 10 | 56 | -34 | 28 | 3.28 | *p* = 0.036 FWE-corrected^*2^ |
| R superior frontal gyrus | 4 | 28 | 66 | 12 | 3.23 | *p* < 0.001 uncorrected |

**^*^** After small volume correction.

^1^ Repeated spatially adjacent peaks. The first peak in order is the uncorrected peak belonging to a cluster with a size of 150, while the second peak in order belongs to a smaller cluster with a size of 26 and replaces the previous one after small volume correction.

^2^ Uncorrected cluster size is 11 and changed to 10 after small volume correction.
