## Supplementary material for "Functional connectivity between the cerebellum and somatosensory areas implements the attenuation of self-generated touch": Table 3-4

**Extended Table 3-4. Peaks that increased their connectivity with the left cerebellum independently of the participants’ behavioral attenuation.** Peaks reflecting greater connectivity with the left cerebellar seed when this is presented in the context of movement (self-generated) compared to when it is presented in the absence of movement (external) (*Movement_0cm_ -by- Touch_0cm_* interaction, Direction: Self > External).

| Brain region | Cluster size (voxels) | MNI coordinates (mm) | | | *z* | *p* |
| --- | --- | --- | --- | --- | --- | --- |
|  |  | x | y | z |  |  |
| L gyrus rectus | 42 | -4 | 26 | -28 | 3.87 | *p* < 0.001 uncorrected |
| R subcentral gyrus | 46 | 62 | -4 | 12 | 3.71 | *p* < 0.001 uncorrected |
| L inferior frontal gyrus, pars triangularis | 9 | -58 | 26 | 12 | 3.61 | *p* < 0.001 uncorrected |
| R central sulcus (M1/S1) | 54 | 58 | -4 | 34 | 3.59 | *p* < 0.001 uncorrected |
| R central sulcus (M1/S1) |  | 50 | -6 | 26 | 3.21 | *p* < 0.001 uncorrected |
| L medial parietal operculum | 38 | -40 | -30 | 16 | 3.54 | *p* < 0.001 uncorrected |
| L parahippocampal gyrus | 16 | -22 | -22 | -26 | 3.46 | *p* < 0.001 uncorrected |
| L hippocampus | 23 | -30 | -34 | -10 | 3.44 | *p* < 0.001 uncorrected |
| L parahippocampal gyrus |  | -22 | -32 | -10 | 3.14 | *p* < 0.001 uncorrected |
| L cerebellum X | 14 | -22 | -30 | -48 | 3.41 | *p* < 0.001 uncorrected |
| L postcentral gyrus | 14 | -56 | -8 | 22 | 3.40 | *p* < 0.001 uncorrected |
| L middle frontal gyrus | 48 | -40 | 14 | 32 | 3.38 | *p* < 0.001 uncorrected |
| L superior temporal gyrus | 10 | -24 | 10 | -46 | 3.36 | *p* < 0.001 uncorrected |
| R superior temporal gyrus | 14 | 70 | -20 | 6 | 3.29 | *p* < 0.001 uncorrected |
| L angular gyrus | 5 | -52 | -74 | 32 | 3.20 | *p* < 0.001 uncorrected |
